# Response time modelling reveals evidence for multiple, distinct sources of moral decision caution

**DOI:** 10.1101/2021.01.27.427353

**Authors:** Milan Andrejević, Joshua P. White, Daniel Feuerriegel, Simon Laham, Stefan Bode

**Author notes:** **Corresponding Author:** Milan Andrejević, Postal Address: Milan Andrejević, Melbourne School of Psychological Sciences, Redmond Barry Building, The University of Melbourne, Parkville, Victoria 3010, Australia.

## Abstract

People are often cautious in delivering moral judgments of others’ behaviours, as falsely accusing others of wrongdoing can be costly for social relationships. Caution might further be present when making judgements in information-dynamic environments, as contextual updates can change our minds. This study investigated the processes with which moral valence and context expectancy drive caution in moral judgements. Across two experiments, participants (*N* = 122) made moral judgements of others’ sharing actions. Prior to judging, participants were informed whether contextual information regarding the deservingness of the recipient would follow. We found that participants slowed their moral judgements when judging negatively valenced actions and when expecting contextual updates. Using a diffusion decision model framework, these changes were explained by shifts in drift rate and decision bias (valence) and boundary setting (context), respectively. These findings demonstrate how moral decision caution can be decomposed into distinct aspects of the unfolding decision process.

## Introduction

Moral judgements are an integral and pervasive part of being human - they undergird our social relationships and form the basis for our legal, political, and governmental institutions^1^. Yet moral judgements are not made in isolation but in a complex informational context; various types of contextual information can influence our moral judgements. For example, moral judgements can be modulated by contextual information regarding the social identities of moral actors and victims^2,3^, their economic class^4^, their relational status^5,6^, the actor’s mitigating circumstances^7^, as well as the victim’s history of moral or immoral actions (i.e. their moral deservingness)^8,9^. Such information can also lead people to change their minds about their moral judgements. Recent research has shown that people flexibly update their judgements upon receiving contextual information, switching from relying on context-independent to context-dependent norms^9^.

In distributive justice scenarios, context-dependent norms are often preferred over context-independent norms. For example, if information regarding individual contributions of actors to shared resources or the history of previous transactions is made available, people prefer splitting resources in accordance with norms that account for such information (e.g., equity norm^10–12^, reciprocity norm^13,14^, indirect reciprocity norm^15,16^), rather than ignoring this information and allocating equal amounts to each individual (equality norm^17–19^). Some individuals prefer context-dependent norms so much that they refrain from making strong judgements prior to the presentation of contextual information^9^. This may reflect the caution of these individuals not to make judgements that may later, upon learning additional information, turn out to be mistaken as they no longer align with preferred context-dependent norms. Thus, caution against selecting a judgment which that is not in line with personal moral norms (i.e. moral decision caution) likely plays an important role in dynamic everyday moral decision-making situations, especially when we are aware that we may learn additional, decision-relevant information in the near future. However, the moral decision process in such situations is poorly understood, partly because there is no adequate process model that would allow investigating various forms of moral decision caution more directly.

Caution has been studied in different areas of decision research, mostly using single-decision tasks involving perceptual and reward-based choices. One form of caution characterized by this research is a tendency for an individual to slow their response time (RT) in order to increase the likelihood of making a correct choice^20^. Generally, participants show this tendency when under explicit instructions to ensure high accuracy^21^, when there are high monetary costs for mistakes^22^, and in conditions of high task difficulty, so as to maximize reward rate^23^. This form of caution enables people to adapt their decision processes to suit changing environmental demands.

This form of caution may also be a useful way for people to adapt their moral decision-making when there is an expectancy of learning more information at the time of the decision. Recent research suggests that people display this form of caution as they learn about the likelihood of outcomes or consequences of their choices. People slow their judgements to reduce the likelihood of errors when they are aware that they are likely to make an error in the given task^24^, as well as when they are aware that the association between their choices and the outcomes resulting from these choices is volatile^25^. Expectancy of learning contextual information in moral judgements also changes the subjective likelihood of judgement errors, defined as choices that appear correct based on the information available at the time of the decision (in line with context-independent norms) but turn out to be incorrect as contextual information is learned (not in line with the preferred context-dependent norms). Therefore, expectancy of learning contextual information may also lead to this form of caution and slow down moral judgements.

However, there is also another form of caution, which is highly relevant for moral judgements: People may particularly slow their RTs when judging someone’s action as morally bad, to increase the likelihood of being correct (according to their personal moral norms) when selecting this option. This tendency can be conceptualised as a *decision bias*, which has been shown to occur in other contexts against choice options associated with smaller rewards^21^, or larger punishments^26^. Morally blaming others is socially risky as it may lead to reprisals if that blame is improperly placed. Indeed, people are more motivated to stay as accurate as possible by ensuring their judgements are up to date with all the available information when making negative judgements^27,28^. However, there is an alternative explanation for why people may take longer when judging someone as bad that is unrelated to caution. Namely, people tend to take longer to evaluate negative information, even when they are not required to make any decisions, and there are no response options to be cautious about. For instance, people report thinking more thoroughly about negative events^29^, they look longer at negative content when scrolling through images^30^, and are longer distracted by morally negative words^31^. Such effects suggest that people take longer to process negatively valenced information^32^. Therefore, there are two distinct explanations for slower RTs when making negative moral judgements: a decision bias (defined as a tendency to be more cautious when judging someone as bad), and a slower rate of evaluation of negative information (i.e. evidence for the negative judgement). For this reason, previous research relying on simple comparisons of mean RTs has been unable to disentangle the cognitive processes underlying the slowing^32^. This is again due to the fact the there is no process model specifically developed for moral decision-making that would allow us to investigate this question.

In other fields of decision science, evidence accumulation models have been widely applied to disentangle parts of the decision process. These process models include mathematically formalized parameters that correspond to evidence accumulation (i.e. the rate at which evidence is evaluated) and the two forms of caution described above, and might therefore be useful for partitioning distinct sources of moral decision caution. One prominent model of this class is the Diffusion Decision Model (DDM)^33–35^ which has been used to study decision-making across a broad range of discrete choice tasks^33,36–39^. The DDM describes the decision process as a continuous accumulation of noisy evidence for different choice outcomes. Once evidence in favour of a particular choice reaches a boundary, a decision is made. These models find substantial support from animal studies where neural firing rates in middle temporal and ventral intraparietal areas found to closely track the trajectory of evidence accumulation^40,41^. Although predominantly used to model perceptual decision-making processes, where sensory evidence is accumulated by the sensory systems, the DDM can be regarded as a universal decision process model, and it has been used to model value-based decisions^42^, sharing and cooperation choices^43–45^ as well as moral decisions^46^. For such higher-level decisions, the accumulation process represents integration of signals from brain areas that calculate subjective value^47^, integrate representations of potential gains and losses^48^, and perform diverse social and moral computations that have not yet been well specified by previous research^44,46^.

The rate of evidence processing (i.e. the evidence strength), and two forms of caution, correspond to specific parameters of the DDM model. In the DDM model, caution against making an error across response options is formalized as the amount of evidence needed to make a choice and is estimated by ***the boundary separation (a)*** parameter. Given the role of this parameter in adapting decision processes to environmental demands^21,24,25^ (as described above), this parameter may increase - corresponding to a wider boundary separation, when expecting contextual information. Biases against one of the response options are formalized as shifts in the starting point of the accumulation process, thus capturing how much people favour a certain response option prior to observing the stimulus, and is estimated by ***the (starting point) bias parameter (z)***. This parameter has been shown to shift towards the response option associated with a reward^21^ and away from response options associated with punishment^26^. In the case of moral judgements, because of the potential social repercussions that come with placing moral blame improperly, the bias parameter may be shifted towards the “good” judgement choice. And third, the average rate of evidence accumulation, capturing the strength of evidence favouring either response option in a task, is estimated by ***the drift rate (v)*** parameter. In visual discrimination tasks, this parameter has been shown to scale with stimulus discriminability^21^. We expected this parameter to scale with the prototypicality of action as morally good (representing adherence to a moral norm) or bad (representing deviation from a moral norm). Moreover, if negative evidence is accumulated slower than positive evidence (independent from potential biased caution against “bad” judgements accounted by the *z* parameter), we expect negative evidence to decrease the drift rate parameter.

In the current study we used a modified version of a recently developed moral judgement task^9,49^ to test the effects of context-expectancy and moral valence on RT and parameters of the decision process, as operationalised by the DDM. We asked participants to observe a variant of the dictator game in which a “Decision-maker” decided to share a proportion of $10 with another person (the “Receiver”; Figure 1a). Participants were aware that these choices were made in a particular informational context. Participants knew either that decision-makers knew nothing about the Receiver, or that they knew how ‘deserving’ the Receiver was, based on their past sharing behaviour towards another person. In each trial participants made moral judgements about the Decision-maker’s sharing action while expecting a contextual update about the Receiver’s deservingness (*context-expectant* condition) or while not expecting an update (*no-context* condition; Figure 1b). In Experiment 1 these two conditions were presented interleaved. Experiment 2 was used to replicate the results using a near-identical paradigm with an independent sample of participants. In the second experiment the two conditions were presented in separate blocks, which further controlled for the possibility that the interleaved presentation of conditions might have had an impact on participants’ decision strategies. Naturally, there are individual differences in the norms people rely on to make such judgements. A majority of people, however, condemn low and endorse high offers^9^. To avoid possible confounding of response times due to potential differences in the reliance on different sets of norms across individuals, and to ensure that the perception of our stimuli as “evidence” for judgement options was roughly consistent across the sample (a necessary assumption of the DDM when fit for a group of individuals in a hierarchical model, see Methods), we limited our investigation in both experiments to this largest subset of participants, who endorsed generosity and condemned selfishness^9^ (implications for limitations will be addressed below).

**Figure 1.**
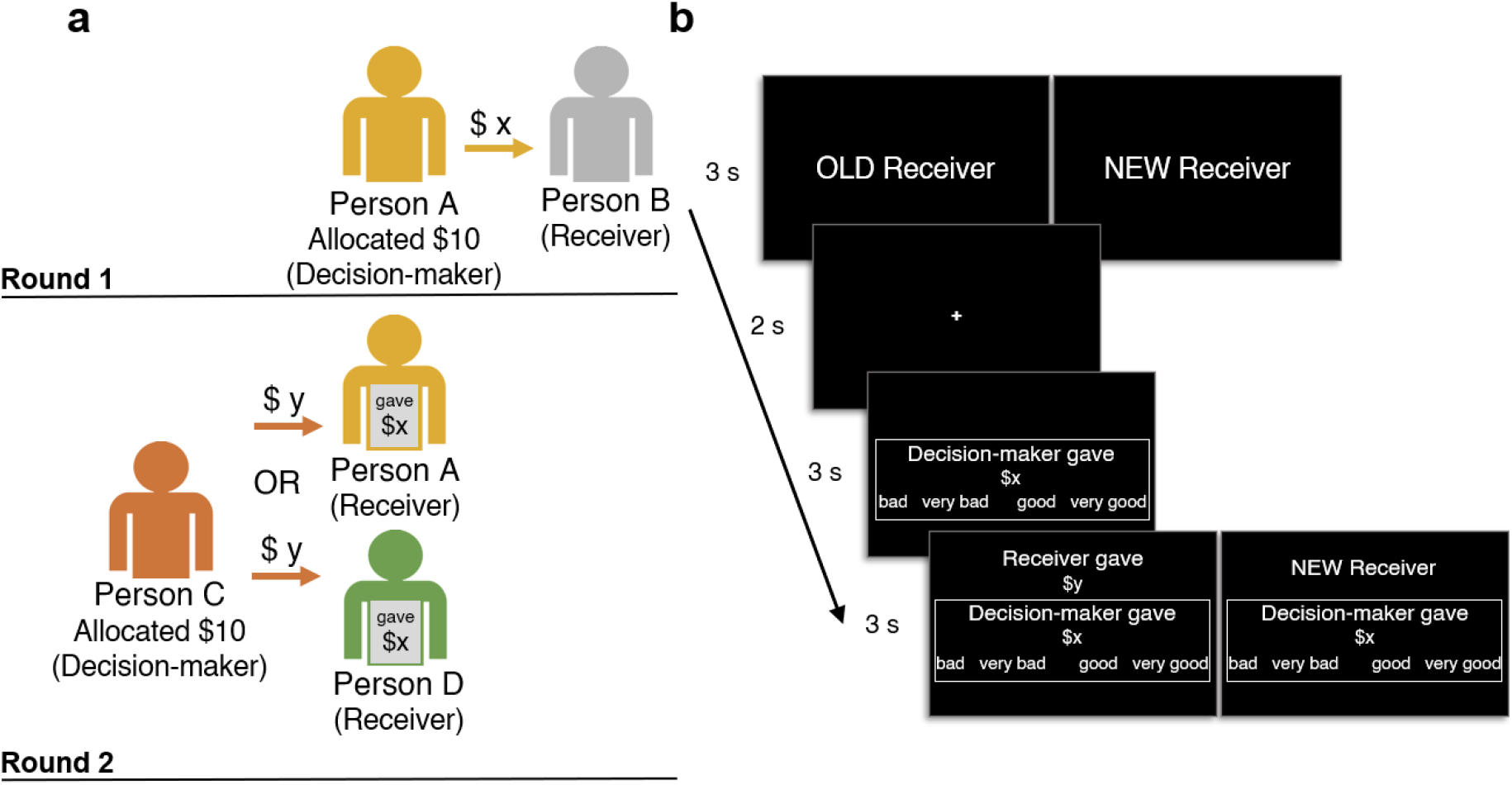
Moral judgement context-expectancy paradigm. (**a**) Depiction of the cover story participants read prior to the experiment about a recently conducted study. The cover-story study was fictitious, but our participants were not informed of this. It involved persons interacting across two rounds: In Round 1, Person A played the role of the Decision-maker and had to decide how to share $10 with their partner, Person B. In Round 2, a new person (Person C) became a Decision-maker and was paired with either Person A or a new person (Person D), and had to decide how to share $10 with their partner. Importantly, Person C knew whether their partner took part in Round 1, and if they did (e.g., Person A), how much they gave when they were the Decision-maker (*$x*). Person C decided to give a certain amount (*$y*) to their partner (either person A or person D, depending on the trial). (**b**) Trial sequence. Participants were presented with information regarding the context-expectancy condition of the current trial. “OLD Receiver” indicated that they would judge the Round 2 Decision-maker who was paired with a Receiver (i.e. Person A) who gave an amount $x to another person in the previous round. Our participants made this judgement without yet knowing this $x amount, but knowing that they would soon learn this information (i.e. the context-expectancy condition); or “NEW Receiver”, indicating that they would judge the Round 2 Decision-maker paired with a new person (i.e. Person D), and that there was no additional contextual information to expect (no-context condition). Next, participants were presented with the amount that the Round 2 Decision-maker gave to their partner and selected their judgement (one of the four options) on a keyboard. After this, in context-expectant condition, the amount that the Receiver had given in the previous round (*$x*) was revealed. In the no-context condition, no additional information was presented. Participants again indicated their judgement of the Decision-maker’s action on their keyboard.

## Results

The selected sample of participants relied on similar norms when performing their judgements (judging low offers as bad and high offers as good) as expected, and there were no systemic differences in the proportions of moral choice for each choice option across expectancy conditions (depicted in Figure 2).

**Figure 2.**
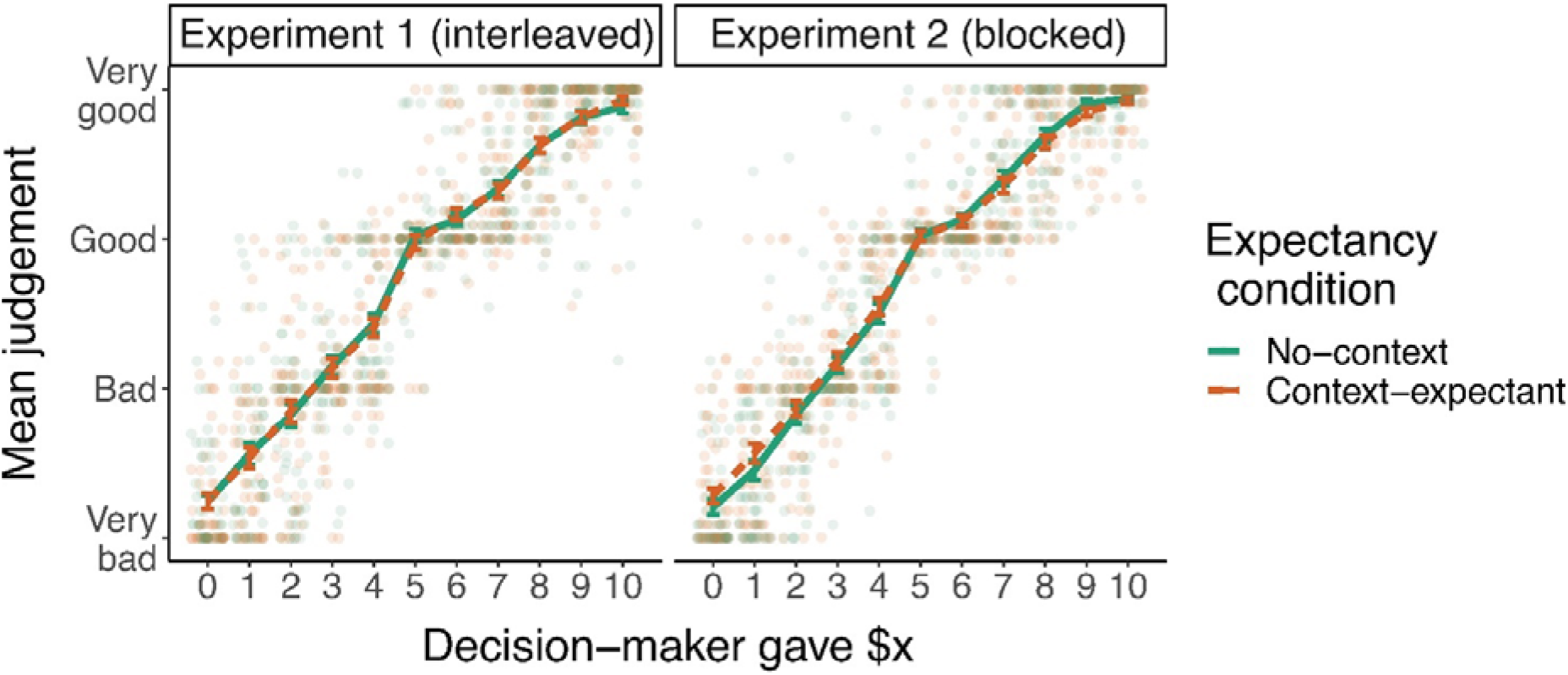
Moral judgement results. Scattered dots indicate mean judgements of Decision-maker’s offer for each participant; lines indicate the mean across participants for context-expectancy (orange) and no-context (green) conditions. Error bars depict the SEM. The monotonic increase in moral endorsement across Decision-maker’s offers indicates that participants condemned low and endorsed high offers. The complete overlap of two the lines representing the conditions indicates that there were no detectable systemic differences in moral judgement across expectancy conditions in the two experiments.

We took two approaches to test for effects of context-expectancy and moral valence on response speed in moral judgements. The first approach was to test for these effects by comparing RTs without formally specifying the decision process. Our predictions for these RT comparisons together with the analysis approach were preregistered (https://aspredicted.org/blind.php?x=dy3qk9). The second approach was to use the DDM to better characterise these effects by comparing model parameter estimates across expectancy and valence conditions.

With regards to our first approach we tested three hypotheses. First, we investigated whether expectancy of contextual information increases caution, by testing whether the RTs of initial judgements were higher in the context-expectant than in the no-context condition. Second, we investigated whether morally negative evidence is evaluated more cautiously and is processed at a slower rate. This hypothesis was operationalised as the assumption that the effect of morally negative valence linearly decreases with the size of the Decision-maker’s offer. We therefore expected a negative relationship between the Decision-maker’s offer and RT. Third, to investigate whether caution when expecting a contextual update is particularly pronounced for negative judgements, we tested whether the slope of the negative relationship between RT and Decision-maker’s offer was steeper in the context-expectant condition. To test these hypotheses, we formulated several Generalised Linear Mixed-effects Models, which included the Decision-maker’s offer, the expectancy condition, and their interaction, as predictors of RT (Supplement 1 Table S2).

The best-fitting model included main effects of Decision-maker’s offer and expectancy condition but did not include an interaction, and the intercepts and the slopes of these two main effects were allowed to vary across individuals (Supplement 1 Table S2). The two main effects were substantial and statistically significant. Context-expectancy led to a 23 ms slowing of RTs, 95% CI [5, 41] in Experiment 1. One possibility is that this effect may have been reduced due to the interleaved design, which could have led to an overspill of decision criteria among trials of different conditions. This was addressed with Experiment 2, which used a blocked design and replicated the context-expectancy effect, which was indeed much larger. Context-expectancy led to a 138 ms slowing, 95% CI [110, 165]. These results support the hypothesis that context-expectancy increases caution. Regarding the second hypothesis, we found that a single dollar reduction in the Decision-maker’s offer predicted a 26 ms slowing of RTs, 95% CI [31, 21] in Experiment 1. This effect again replicated in Experiment 2 showing 28 ms slowing for each dollar reduction in the Decision-maker’s offer, 95% CI [31, 24]. These results support the hypothesis that that negatively valenced evidence is evaluated more cautiously or takes longer to process. The two main effects were consistent across quantiles (RT quantiles by condition are displayed in Figure 3). They were also robust across different models (Supplement 1 Table S2) and across alternative approaches to modelling RT distributions (Supplement 1 Table S4 and S5). As for the interaction effect, there was no evidence across these two studies supporting the hypothesis that effects of negative valence are more pronounced when people are expecting a contextual update. Models including the interaction effect had poorer fits, and confidence intervals for this interaction parameter consistently included zero.

**Figure 3.**
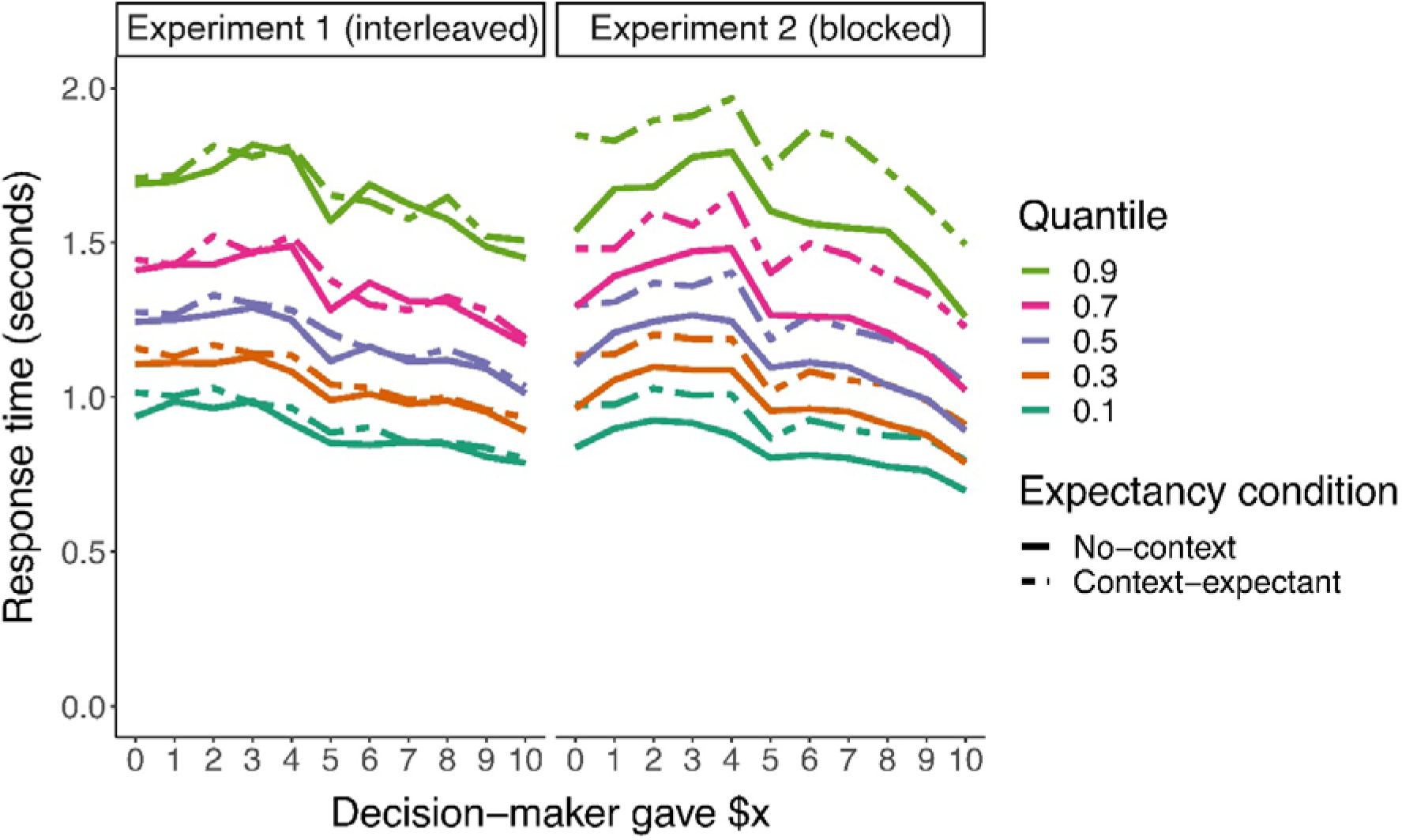
Moral Judgement RT quantiles across Decision-maker offers and expectancy conditions. The graph shows group mean quantile values across participants. The general pattern of results was consistent across quantiles and across two studies: there was a slight increase in speed for higher Decision-maker offers; and there was a slight slowing in context-expectancy trials in Experiment 1 (dashed lines higher than solid lines), which was more pronounced in Experiment 2.

Next, to better characterise these patterns of RT effects, and to test our predictions regarding the relationships between context-expectancy, moral valence and components of the decision process, we fitted a Diffusion Decision Model. To test whether context-expectancy increased the general amount of caution across judgement options (i.e. boundary separation), we computed two *a* parameters, one for each expectancy condition, and compared them. We expected: *a*_*context-expectant*_ > *a*_*no-context*_. To test whether moral prototypicality of Decision-maker’s offers reflected stronger evidence for judgement options (with lower offer magnitude reflecting evidence for “bad” option, and higher offers reflecting stronger evidence for the “good” option, in line with the norms applied by the selected sample), we fitted a *v* parameter separately for each Decision-maker’s offer. The *v* parameter was signed, meaning negative values indicated evidence for “bad” judgement and positive values indicated evidence for “good” judgement. We tested for a monotonic positive relationship between the offer magnitude and the *v* parameter. Moreover, to test whether negatively valenced evidence is accumulated more slowly than positively valenced evidence, we tested whether the estimates of the *v* parameter were in absolute terms (drift towards either “good” or “bad”) larger for high as opposed to low Decision-maker’s offers. We expected: |*v*_*0-4*_| < |*v*_*6-10*_|. Finally, to test whether participants were more cautious against making “bad” judgements, independent of the tendency to more slowly accumulate negatively valenced information, we tested whether the *z* parameter differed from .5 (which would indicate no starting point bias), and whether the *z* parameter was biased in the direction of ‘morally good’ judgement. The position of decision bounds with respect to the starting point were standardized as 1 for ‘morally good’, and 0 for ‘morally bad’ judgements, hence we expected *z* > .5.

First, we formulated a hypothesised model (***m1***), which included separate *a* parameters for each expectancy condition, separate *v* parameters for every offer value, and a *z* parameter. We then tested whether the use of this model, which allowed us to test our specific hypotheses, was justifiable and appropriately explained our data, by comparing it to a null model (***m0***), which did not include differences between conditions for any parameter. We used the Deviance Information Criterion (DIC) to compare the model fits (lower value indicates better fit)^50^. We found that this model provided a substantially better fit to the data (Experiment 1 DIC = 13396.303; Experiment 2 DIC = 15021.171) than the null model (***m0***) (Experiment 1 DIC = 29569.801; Experiment 2 DIC = 30071.387). Additionally, we ran Posterior Predictive Check (PPC) simulations of the two models. Simulated data from model ***m1*** more closely resembled the quantile structure of the observed RT data (Supplement 2 Figure S10). The ***m1*** model simulation also reproduced the observed rates of judgements across Decision-maker offers (Supplement 2 Figure S8) and patterns of changes in RT distributions across different Decision-maker’s offers and expectancy conditions (Supplement 2 Figure S9), and overall provided an excellent fit to the data.

Next, we tested for hypothesised differences in the ***m1*** model parameters across conditions. Statistical significance was defined as the posterior probability for the hypothesised difference exceeding .95. Consistent with our hypothesis that context-expectancy increases caution against making errors, the *a* parameter estimate was nominally larger in the context-expectant condition compared to the no-context condition; however, this difference was not statistically significant in Experiment 1 (posterior P(*a*_*context-expectant*_ > *a*_*no-context*_) = 0.913) (Figure 4a). In Experiment 2 this difference was statistically significant (posterior P(*a*_*context-expectant*_ > *a*_*no-context*_) > 0.999, Figure 4b). As for the drift rate (*v)*, we expected this parameter to monotonically increase with the value of the Decision-maker’s offer. We observed a perfect monotonic relationship across both experiments (see Figure 4c and d). To test our hypotheses regarding the reduction in absolute drift rate when processing negative moral valence as compared to positive valence, we compared the *v* parameter for negative stimuli (Decision-maker gave $0-4) with positive stimuli (Decision-maker gave $6-10). Consistent with our hypothesis we found a large and statistically significant decrease in absolute drift-rate for negative stimuli (Experiment 1 posterior P(|*v*_*0-4*_| < |*v*_*6-10*_|) > 0.999; Experiment 2 posterior P(|*v_0-4_*| < |*v*_*6-10*_|) > 0.999, see Figure 4c and d). To ensure that this effect was not due to a perception of Decision-maker’s offer of $4 as neutral as opposed to negative, we repeated these analyses on a more constrained set of stimuli by excluding offers $4 and $6, and the effect survived in both studies (Experiment 1 posterior P(|*v*_0-3_| < |*v*_*7-10*_|) = 0.999; Experiment 1 posterior P(|*v_0-3_*| < |*v_7-10_*|) = 0.999). To test our hypothesis regarding the shift of the bias parameter (*z*) away from the ‘bad’ and toward the ‘good’ judgement option, we tested whether the *z* parameter was larger than .5. Consistent with our hypothesis we found estimates of *z* parameter to be larger than .5 in both studies (Experiment 1 posterior P(*z* > 0.5) > 0.999; Experiment 2 posterior P(*z* > 0.5) > 0.999).

**Figure 4.**
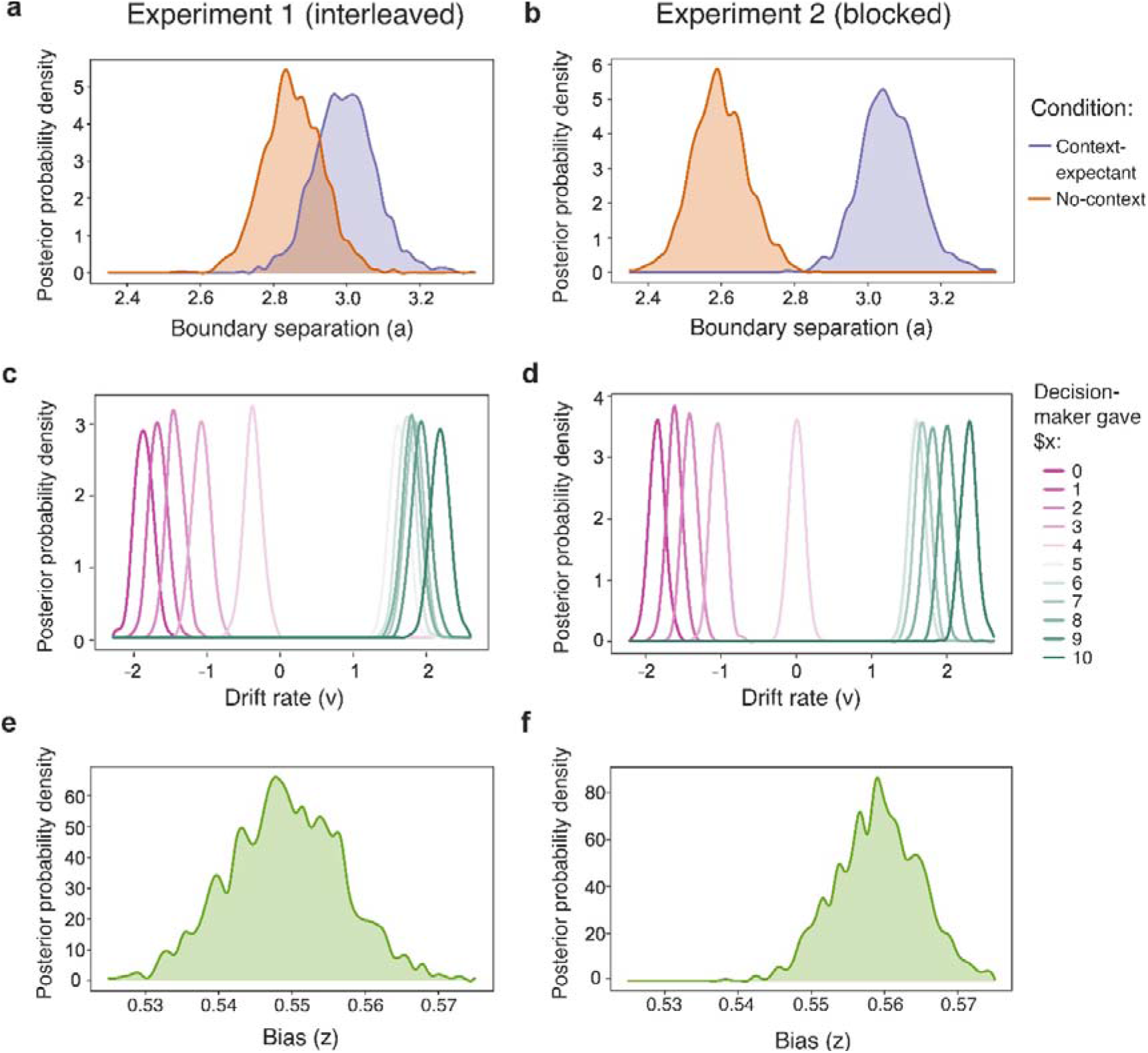
Bayesian posterior probability distributions for Diffusion Decision Model parameters *a, v* and *z* for both Experiments. (**a**) In Experiment 1, the boundary separation parameter (*a*) estimate, although overlapping, was slightly higher for the context-expectant condition, which is in line with the hypothesis that context-expectancy increases caution. (**b**) In Experiment 2, this difference was replicated with a larger effect and there was minimal overlap between the two posterior distributions. (**c**) In Experiment 1, the drift rate parameter (*v*) monotonically increased with higher Decision-maker’s offers, suggesting that higher offer numbers provide more evidence for the judgement option ‘good’ and less for ‘bad’. Positively valenced actions (DM gave more than 6) had higher absolute drift rates towards option ‘good’ than negatively valenced actions did towards option ‘bad’ (DM gave less than $4), which suggests that participants processed negatively valenced actions slower than positively valenced actions. (**d**) These effects replicated in Experiment 2. (**e**) In Experiment 1, participants showed a bias towards judging ‘good’ (*z* parameter >.5), which is in line with the hypothesis that people may be more cautious when making negative judgements. (**f**) This effect replicated in Experiment 2.

## Discussion

We investigated the effects of context-expectancy and negative moral valence on moral decision caution in third-party moral judgements of sharing actions. Both factors were hypothesised to slow moral judgements, albeit impacting different aspects of the decision-making process. Specifically, we examined these effects by comparing RTs and parameter values of the DDM across judgements of fairness-related actions (i.e. offers of different magnitudes) as well as context-expectancy conditions. Our results show a significant slowing of RT in the context-expectancy conditions, as well as for morally negative actions; however, there was no interaction between the two factors. Moreover, these effects were well accounted for by differences in multiple DDM parameters. The boundary separation parameter was larger in the context-expectancy condition compared to the no-context condition, pointing to more caution to avoid erroneous responses (across judgement options) in the former condition. In addition, signed drift rates increased with the Decision-maker’s offer, suggesting that lower offers corresponded to stronger evidence for negative judgments and higher offers corresponded to stronger evidence for positive judgements. Absolute drift rates were smaller for negatively valenced offers, supporting the notion that negative evidence is accumulated at a slower rate than positive evidence, for reasons most likely not related to moral decision caution per se. Additionally, the starting point parameter showed a bias against “bad” judgements, suggesting that people also slowed their negative judgements as they were particularly cautious about them.

Our findings that participants slowed their judgments when expecting contextual information is consistent with previous research showing that people are more cautious when aware that they are more prone to making mistakes^24,25^. Notably, previous research has demonstrated this effect for decision mistakes in tasks in which people are not given additional information or a chance to change their minds^24,25^. The current findings show that this effect also extends to dynamic decision-making contexts, in which learning additional information can lead to changes of mind. Crucially, here we show that this type of caution can be explained by the widening of the decision boundary separation in a process model of decision-making.

Finding that the expectancy of contextual information increases the boundary separation also highlights the importance of contextual information for moral judgements. This finding is consistent with previous research that showed that contextual information influences the judgements that we make^2–8^, and that some people make less extreme good/bad judgements when expecting contextual information^9^. To note, we did not find an adjustment of the judgement itself (see Figure 2), but the relatively course four-point scale might not have been ideal to capture any potential subtle effects that might have occurred but could not be expressed without a finer scale. The difference in response times, however, was observed even though the expected contextual information could never directly impact the initial judgment. This is important because it shows that context-dependent norms affect our judgements even when contextual information is not yet known, a point which has been overlooked in the moral judgement literature.

We further found that participants were slower when evaluating lower offers, which is in line with both the idea that people take longer to process negative evidence^29–32^, as well as with the idea people are more cautious against judging people as bad, as negative judgements have higher social repercussions for individuals^27,28^. Our DDM results further support each of these accounts separately. Firstly, our finding that the drift rate was slower for lower offers as compared to higher offers is in line with the idea that people accumulate negative evidence at a slower rate^29–32^. Secondly, we found that participants showed biases, or caution, against judging moral actions as bad, independent of taking longer to process negative evidence. Previous research on financial decision-making showed similar bias parameter shifts away from options associated with less favourable monetary outcomes^21–23,26^. Our results extend these findings to moral judgement valence, suggesting that people are inclined to default to positive judgements. This may be because of the sensitivity of the bias parameter to social outcomes, such as the repercussions that come with placing moral blame improperly^27,28^. Overall, our findings suggest that people take longer to make judgements about negative actions both because it takes them longer to process negative information, and because they favour positive judgements.

Our finding that people have biased caution against making negative judgements complements recent findings showing that people are more prone to adjust and change negative rather than positive beliefs about others^28^. Although negative beliefs are more susceptible to change, our results suggest that people are more cautious to form these beliefs in the first place. Together, these findings suggest that people are more careful about being accurate when evaluating morally negative evidence, both in terms of changing their minds when receiving information updates^9,28^, and by allowing themselves time to consider all the information that is available when prompted to make a judgement.

Our finding that signed drift rates showed a monotonic relationship with the magnitude of Decision-makers’ offers is in line with the idea that moral prototypicality of the action determines the quality of evidence for moral badness and goodness. Previous research showed that drift rate scales with perceptual discriminability of the stimuli in classical perceptual decision tasks^21^. Our findings suggest that this effect generalizes to moral decisions, which is in line with the idea that moral prototypicality (i.e. how well a moral action represents adherence to or deviation from a moral norm) equates to moral discriminability and determines the rate of moral decision evidence accumulation.

We did not find support for our hypothesis that context-expectancy would interact with the moral valence effect. Our RT results instead suggest that these two effects were additive. These results are somewhat in discord with a previous finding that some participants reduced the intensity of their negative moral judgements (but not positive moral judgements) when expecting a contextual update^9^. There are several explanations for this discrepancy. This previous finding may be specific to moral judgements reported on a continuous scale. It may also occur only in smaller subset of people. Our strict focus on a subsample of people that condemn low offers may have excluded the people that reduce the intensity of this condemnation and show this effect. Future studies could preselect samples of people who show this effect and characterise their decision process specifically.

There are several remaining open questions that should be investigated in future studies. One outstanding question is whether the DDM can be applied to better characterise aspects of moral decision-making across a wider range of contexts. While the DDM has primarily been used to derive psychologically meaningful parameters in perceptual decision tasks^33,36,37,42^, and has only been applied to a small range of social and more specifically moral tasks^44–46,51,52^, our results illustrate that the DDM can be a powerful tool for dissociating parts of the decision-making process in social tasks. Our findings show that the DDM can be used to clearly partition RT variance in such tasks, and the consistency of results across two samples suggest that this partitioning is reliable. Future studies could test how well our findings generalize to other kinds of judgement tasks (e.g., traditional moral dilemmas), other moral norms (e.g., concerning harm), and other kinds of contextual information (e.g., relational status between moral actors). It could further be tested whether there is an even better model within the DDM framework to capture the process of moral judgement. We have restricted our analyses to the most plausible (and hypothesis driven) model instead of exploring the full space of all possible models, which was beyond the scope of our study. Future research, however, can extend this framework, for example by including parameters such as collapsing decision bounds^42,53,54^, or by allowing for inter-trial variability of some parameters^55,56^ to further improve the model fit; however additional theoretical work is needed to justify inclusion of such variations for the current context. Additionally, our study remains agnostic to neural mechanisms behind the moral decision process. To better understand the computation behind moral decisions, future studies should investigate the neural correlates of these computations.

To conclude, our findings identify expectancy of learning new contextual information and moral valence as impacting two distinct forms of moral decision caution. While context expectancy slows moral judgements to reduce erroneous responding in general, negatively valenced information also leads to slower judgements, presumably reducing the likelihood of making an erroneous negative judgement. Additionally, we also show that this effect of negative valence occurs in addition to another effect – that negative evidence is accumulated at a slower rate than positive evidence. These findings improve our understanding of processes underlying moral decision-making in dynamic situations and provide a foundation for future research on neural mechanisms underlying moral decisions.

## Materials and Methods

### Participants

The study was approved by the Human Research Ethics Committee of the Melbourne School of Psychological Sciences (Ethics ID 1750046.3). Participants were compensated with course credit or monetary remuneration ($15). Participants were right-handed, fluent in English, and had normal or corrected-to-normal vision.

For Experiment 1 (interleaved design), 77 people participated (50 female, 27 male, *M*_*age*_ = 24.70, *SD* = 7.40, range: 18–69 years). Eleven participants were excluded from the sample for data quality reasons: nine participants failed an attention-check (i.e. had given incorrect answers in more than 40% of catch trials of either category; see below), and two participants had missing responses for over 5% of trials, again suggesting a lack of attention. We preselected the final sample such that all included participants would rely on the same moral norms to make their judgements. This was done to avoid possible confounding of response times due to potential differences in norm-related information processing across norms, and to ensure that all participants were assigning moral meaning to presented stimuli in a similar manner (which is a necessary assumption of the DDM when fit for a group of participant datasets). Based on previous research using a similar task, we expected the largest group to be participants who endorsed high and condemned low offers^9^. A strong positive correlation between moral judgements and Decision-maker’s offer was typical for this largest group. We excluded eleven participants who did not show this strong positive correlation (Spearman correlation was below *r* = .5). All of these criteria were predefined and preregistered (http://aspredicted.org/blind.php?x=n2fi7g). The final sample consisted of 55 participants (37 female, 18 male, *M*_*age*_ = 24.84, *SD* = 5.86, range: 18–43 years).

For Experiment 2 (blocked design), 76 members of the University of Melbourne community were recruited (47 female, 28 male, 1 other, *M*_*age*_ = 24.29, *SD* = 3.77, range: 19–39 years). Nine participants were excluded to ensure data quality: six participants failed the attention-check criterion (see above) and three had missing responses for over 5% of trials. Another ten participants were excluded because their moral judgements did not correlate strongly with the Decision-maker’s offer (Spearman correlation *r* < .6). All of these criteria were predefined and preregistered (https://aspredicted.org/blind.php?x=dy3qk9). The final sample consisted of 57 participants (38 female, 18 male, 1 other, *M*_*age*_ = 24.34, *SD* = 3.80, range: 20–39 years).

#### Apparatus

The experimental task was programmed in MATLAB (MathWorks, version R2015b) and presented using PsychToolbox-3^57^. Participants sat at a viewing distance of approximately 80 cm from the monitor (ASUS ROG Swift PQ258Q 24.5” HD with a 60 Hz screen refresh rate). The experiment was conducted in a well-lit solitary room. Participants made responses on a black Hewlett-Packard KU1469 QWERTY keyboard. The “z”, “x”, “.” and “/” keys were covered with white stickers to indicate to participants that these were the primary buttons to be used in the experiment. They were instructed to place their fingers on these keys in preparation for every trial in the following manner: the middle finger and the index finger of their left hand were to be placed on the “z” and “x” keys, respectively, and the index finger and the middle finger of their right hand were to be placed on the “.” and “/” keys, respectively.

### Experimental Paradigm

#### Cover Story

Participants first read a cover story about a recently conducted experiment investigating people’s economic decisions. This experiment was fictional, but participants were not informed of this. In the fictional experiment a group of people, assigned to pairs, completed a two-round variant of the dictator game (for the original dictator game, see ref ^58^). In the first round, one person (the “Decision-maker”) in each pair was given $10 and decided how much thereof to share with their partner, the “Receiver”, to whom they could give any whole dollar portion (i.e. any amount $0–$10). In the second round, the same task was repeated except with people taking new roles — first round Decision-makers became Receivers in the second round — and were assigned different partners. Some of these new partners were Decision-makers in the first round of the experiment (“Old Receivers”) and some of them were not (“New Receivers”). Importantly, second round Decision-makers were aware whether their partner was an Old Receiver or a New Receiver. If their partner was an Old Receiver, they were also aware how much money their partner had shared with another person in the first round of the experiment. A visualisation of this cover story is shown in Figure 1a.

#### Instructions

This cover story along with the description of the experimental task were presented to participants via text interleaved with animated depictions. Participants read the instructions and attended to animations at their own pace. Participants were then required to pass, with 100% accuracy, a test comprised of 32 true–false questions which assessed their understanding of both the cover story and the experiment instructions. Participants could attempt this instruction-check test three times. If they experienced troubles completing the quiz, participants could return to the cover story or instruction presentations to clarify their understanding or ask questions of the experimenters for the same. Participants were required to pass this test before continuing to the experiment.

#### Experimental task

Participants were asked to observe a series of independent transactions that various Decision-makers made towards various Receivers as described in the cover story. Each trial started with the participant being shown, for 3 s, whether the Receiver for that trial was an “OLD Receiver” (for context-expectant trials) or a “NEW Receiver” (for no-context trials) which corresponded to whether the Receiver participated in the first round of the fictitious experiment. Then, a fixation cross was presented in the middle of the screen for 2 s. Participants were then presented with the phrase “Decision-maker gave: $y” where y was an integer from the set Y = {0, 1, 2, … 10} (“Decision-maker offer”). Simultaneously, response options “very bad”, “bad”, “good” and “very good” were presented below the Decision-maker offer. Participants selected their response, with a maximum response window of 3 s, to indicate how morally good or bad they believed this Decision-maker’s action was by pressing the button on the keyboard corresponding to the position of the presented option. To control for possible RT differences that could arise due to differences in motor execution across different fingers, participants were randomly assigned one of four possible mappings of responses to buttons, and this mapping remained the same throughout the experiment. Four mappings were selected to ensure that across participants any of the four fingers was mapped onto each response option. For consistency, none of the mappings had a monotonically increasing or decreasing order in space.

Once participants made their response, the corresponding response option immediately changed colour to yellow until the 3 s time-limit had elapsed (or for 0.3 s if the judgment was made between 2.7 s and 3 s) before reverting to white. This was done to assure participants that their response had been recorded.

Participants were then shown another fixation cross above this information for 0.5 s. The stimuli presented next differed depending on the experimental condition of the trial. In context-expectant trials, participants were presented with the phrase “Receiver gave: $x”, where x was an integer from the set X = {0, 1, 2, … 10}, providing the contextual information of how much the Receiver had given when they were a Decision-maker the first round. In the no-context trials, participants were presented with the phrase “NEW Receiver”, reminding them that the Receiver had not participated in the first round, and thus there was no contextual information about them available. In both conditions, participants made a second moral judgment, within 3 s, about the Decision-maker’s action (not the Receiver’s prior action). Once this response had been made, the corresponding response option changed colour to yellow until the 3 s time-limit had elapsed (or for 0.3 s if the judgment was made between 2.7 s and 3 s), after which a new trial began.

There were 121 trials in each condition, totalling 242 trials per participant. This was chosen such that in the context-expectant condition, participants made moral judgments about all possible combinations of the Decision-maker’s offer (i.e. Decision-maker gave $0–$10) and the Receiver’s prior offer (Receiver gave $0–$10; 11 × 11 = 121). To ensure there was symmetry between the experimental conditions, we also included 121 trials for the no-context condition. In Experiment 1 the order of these 242 trials was randomised for each participant and the two trial types alternated randomly (i.e. the two conditions were interleaved). In Experiment 2, we used a version of the experiment with the two expectancy conditions presented in separate blocks. There were 40-41 trials of the same kind in each block and 6 alternating blocks in total. The order of trials was randomised for each participant, and the participants were randomly assigned one of the two alternating block sequences.

##### Questionnaires

Following the experiment, participants completed various personality measures. We administered the agreeableness section of the HEXACO Personality Inventory-Revised (HEXACO)^59^, a brief set of self-report measures for political orientation^60^, the Social Dominance Orientation scale (SDO)^61^, the Consequentialist Thinking Scale (CTS)^62^, and basic demographic measures. We will analyse and report the questionnaire results in a separate publication.

##### Experiment Feedback and Instruction Checks

Participants were instructed to respond as quickly and accurately as possible and always give a response. If they failed to do so within the 3 s time limit, they were presented with feedback at the end of that trial advising which response was missing (or both) and to “please make sure you always respond”. Two types of attention-checks were also dispersed throughout the experiment. In one, participants were required to report the values seen in the current trial; that is, the amounts that the Decision-maker and/or the Receiver had given. Participants responded by entering this value into number keys on the keyboard. For the second attention-check participants had to report, via button press, whether the Receiver in the current trial was an Old Receiver or New Receiver. Participants were instructed that both these attention-check trials would occur at random times during the experiment.

### Statistical Analyses

#### Regression Analysis

RTs for the first moral judgement were modelled with the Generalised Linear Mixed Models (GLMMs) approach which is a form regression suitable for hierarchical data (e.g. data of multiple individuals in several conditions) that is not normally distributed. Invalid trials (i.e. trials without any response) were excluded from all the analyses (0.72% of all trials in Experiment 1, and 1.17% in Experiment 2). GLMMs are superior to the common practice of transforming data before applying an ordinary-least-square linear mixed model^63^. GLMMs were specified as follows: An identity link was used because it assumes that RTs are direct measures of the duration of the decision process, rather than functional transformations of this duration^63^. A gamma distribution was used as the conditional distribution as it provided a good empirical fit to the data. Moreover, gamma-distributed GLMMs have been used in numerous RT studies with similar tasks^64–67^. Lastly, random effects were included in the model to account for individual differences.

We compared a list of theoretically plausible candidate models which were derived with an increasingly complex random effects structure, as shown in Supplement 1 Table S1. For each random effect structure, a model was fit both with and without a fixed interaction parameter. For all models, the random effects were allowed to correlate; that is, the model had an unstructured variance-covariance matrix. Model parameters were estimated using maximum likelihood estimation via the Laplace approximation, implemented with the *glmmTMB* package^68^ in the R statistical programming environment (version 3.6.1). We selected the best fitting model using the Akaike Information Criterion (AIC). AIC was preferred over the likelihood ratio test, because not all compared models were nested, and because, unlike the likelihood ratio test, the AIC method helps prevent overfitting^69^. AIC was also preferred over the Bayesian Information Criterion^70^ because it was unlikely that any of our candidate models are the true model, which better agreed with the assumptions of AIC^71^. Akaike weights^72^ were calculated for all candidate models as a means to quantify the relative merits of the competing models, and the degree to which one model should be preferred over the others. Confidence intervals (and where necessary, *p* values) for fixed effects were calculated for most models using Wald’s *z* method^73^. The fixed parameter effects from the best fitting model, and their 95% confidence intervals, were then used to test our hypotheses.

#### Diffusion Decision Model Fitting

Participants’ RT and decision data were fit in the Python 3.6 programming environment on a High Performance Computing Cluster^74^, using the Hierarchical Drift Diffusion Model (HDDM) package^75^. This package implements a hierarchical Bayesian Markov Chain Monte Carlo (MCMC) estimation of the DDM with four free parameters (*a*, *v*, *z*, and *t*). HDDM estimates these parameters for each individual, as well as at the group level (which are the estimates we report in this publication). This analysis was not preregistered, but was run separately for Experiment 1 and Experiment 2 samples allowing us to assess whether the findings replicated across samples. Estimation procedure implemented in the HDDM package was chosen as it outperforms other estimation techniques and can accurately recover model parameters based on a small number of observations per participant, especially for participant sample sizes larger than 20^75^. Since the DDM is sensitive to outliers, it is recommended to devise exclusion criteria that ensure that some of the contaminant RTs are excluded whilst ensuring that criteria do not exclude larger portions of the data (e.g., more than 1%)^76^. We conservatively excluded trials in which reaction time was faster than 0.2 s (0.05% of valid trials in Experiment 1 and 0.27% of valid trials in Experiment 2), and slower than 2.8 s (0.37% of valid trials in Experiment 1 and 0.37% of valid trials in Experiment 2). The DDM was designed for binary decisions (e.g., “good” versus “bad”), which means that in order to model our data using the DDM, we simplified our data by collapsing across “very good” and “good” responses (*good* judgement) and across “very bad” and “bad” responses (*bad* judgement). We formulated two models to address our hypotheses: ***m0*** – the null model which assumes no difference between conditions when estimating DDM parameters; ***m1* –** the hypothesised model, in which parameter *a* was allowed to vary across two expectancy conditions (*a*_*context-expectant*_ and *a*_*no-context*_), and parameter *v* was allowed to vary across the range of values of Decision-Maker’s offers (*v_0-11_*). For our Bayesian parameter estimation we used the default non-informative priors in the HDDM package^75^. This is the recommended option for novel tasks that are substantially different from typical perceptual decision-making paradigms prominent in the DDM literature^75^. We obtained parameter estimates by generating a chain of 2500 MCMC samples of the joint Bayesian probability posterior distributions of all parameters at both participant and population level, and discarding the first 500 samples (as recommended in ref ^75^). We evaluated chain convergence using Gelman-Rubin diagnostic over five repeated chains (R□<1.1 for all parameters and at all levels across Experiment 1 and Experiment 2). The two models – our theoretically plausible ***m1*** and the null model ***m0*** – were compared using the Deviance Information Criterion (DIC) goodness of fit measure, which penalises for model complexity. Additionally, we also assessed goodness of fit by performing the posterior predictive check procedure, by which we generated simulated data based on posteriors estimates and compared it to empirically observed data (Supplement 2 Figures S8–10). After establishing that the m1 model outperformed the null model and provided an excellent fit for our data, we tested our specific hypothesis regarding *a, v* and *z* parameters by directly comparing the Bayesian probability posteriors generated by the above-described MCMC procedure.

## Data Availability

Data of all participants, materials including the instructions and the task code, as well as the analyses scripts that support the findings of this study are publicly available on an Open Science Framework (OSF) repository (DOI: 10.17605/OSF.IO/EPD63).

## Acknowledgements

We thank Pragya Arora for her help with data collection as well as Gabriel Ong and William Turner for helpful discussions.

## Author Information

### Contributions

M.A., S.B., and D.F. contributed to conception and design. M.A. programmed the experiment. M.A. and J.W. collected and analysed data. M.A. drafted the article. All authors reviewed and revised the manuscript.

## Supplement 1: Regression Analysis (GLMMs)

### Model Fitting

Comparison of AICs across six candidate models (described in Table S1) in Experiment 1 showed the best model fit for model 5 (see Table S2). This was replicated in Experiment 2. AIC weights for the winning model indicated that the probability that it is the best model of the whole candidate set is very high (95.1% chance for Experiment 1 and 99.9% chance for Experiment 2). These model comparisons suggest there was no interaction effect in the data. Further, even in models that included the interaction as a fixed effect (e.g., models 2 and 4), this effect was not significant across two samples. Finally, the Nakagawa conditional *R*^2^ for the winning model indicates random and fixed predictor variables explain a high portion of variance in the model (35.7% in Experiment 1 and 31.8% in Experiment 2). Nakagawa marginal *R*^2^ indicates that a notable portion of model variance is explained by the fixed predictor variables (4.9% in Experiment 1 and 7.9% in Experiment 2).

The winning model can be mathematically described as follows:

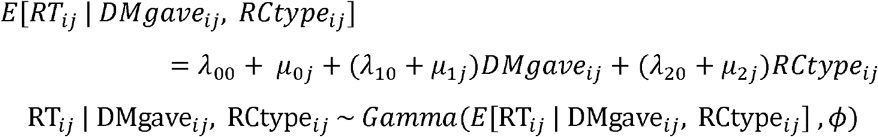

where:

*i* = trial

*j* = participant

*ϕ* = dispersion parameter, which remains constant for different *E*[*Rr*_*ij*_ *DMgave*_*ij*_, *RCtype*_*ij*_]

*μ*_0j_ ~ N(0, σ_0_^2^)

*μ*_1j_ ~ N(0, σ_1_^2^)

*μ*_2j_ ~ N(0, σ_2_^2^)

That is, the best fitting model had random participant-level effects for the intercept (*μ*_0j_), DMgave (*μ*_1j_) and RCtype (*μ*_2j_); and fixed trial-level effects for the intercept (*λ*_00_), DMgave (*λ*_10_) and RCtype (*λ*_20_). Analysis of the fixed effects estimates shows that there was a significant negative effect for Decision-maker offer and a significant positive effect for context-expectancy (Table S2). Table S3 lists the random-effects structure of the model. Other models were also evaluated and compared to ensure that the parameter estimates in this model were robust to assumption violations, and not a mere artefact of our choice of conditional distribution and link function (i.e. a gamma conditional distribution and identity link function). The fixed effect parameters and their standard errors are approximately equal when a log link function was used (Table S4). Similarly, even using a gaussian conditional distribution (i.e. an ordinary linear mixed model) yields similar fixed effects results (Table S5). These analyses showed that results remain similar regardless of the exact methodology utilised.

**Table S1.** Fixed and random effects included in each candidate generalised linear mixed model.

| Candidate Model | Random effects included |  |  | Fixed effects included |  |  |  |
| --- | --- | --- | --- | --- | --- | --- | --- |
|  | Intercept | DM offer | Expectancy Condition | Intercept | DM offer | Expectancy Condition | Interaction |
| 1 | x |  |  | x | x | x |  |
| 2 | x |  |  | x | x | x | x |
| 3 | x | x |  | x | x | x |  |
| 4 | x | x |  | x | x | x | x |
| 5 | x | x | x | x | x | x |  |
| 6 | x | x | x | x | x | x | x |

*Note.* DM = Decision-maker.

**Table S2.** Model comparison table of all candidate models across two studies (interleaved and blocked).

| Experiment | Candidate Model | Random effects included | Fixed effects (with 95% Confidence Interval) |  |  |  | Model fit statistics |  | Nagakawa R<sup>2</sup> |  |
| --- | --- | --- | --- | --- | --- | --- | --- | --- | --- | --- |
|  |  |  | Intercept | DM offer | RC type | Interaction | AIC | AICw | Conditional | Marginal |
| 1 | 1 | Intercept | 1.363*<br>[1.309, 1.418] | -0.025*<br>[-0.027, -0.023] | 0.025*<br>[0.013, 0.037] | NA | 10769.96 | .000 | .335 | .047 |
| 1 | 2 | Intercept | 1.359*<br>[1.304, 1.415] | -0.024*<br>[-0.027, -0.022] | 0.033*<br>[0.009, 0.058] | -0.001<br>[-0.005, 0.002] | 10771.38 | .000 | .335 | .047 |
| 1 | 3 | Intercept and DM offer | 1.366*<br>[1.307, 1.425] | -0.026*<br>[-0.031, -0.021] | 0.025*<br>[0.013, 0.037] | NA | 10600.50 | .033 | .355 | .048 |
| 1 | 4 | Intercept and DM offer | 1.363*<br>[1.303, 1.422] | -0.025*<br>[-0.030, -0.020] | 0.032*<br>[0.007, 0.056] | -0.001<br>[-0.005, 0.003] | 10602.12 | .015 | .355 | .049 |
| **1** | **5** | **Intercept, DM offer, and RC type** | **1.367\***<br>**[1.307, 1.427]** | **-0.026\***<br>**[-0.031, -0.021]** | **0.023\***<br>**[0.005, 0.041]** | **NA** | **10593.80** | **.951** | **.357** | **.049** |
| 1 | 6 | Intercept, DM offer, and RC type | 1.363*<br>[1.303, 1.424] | -0.025*<br>[-0.030, -0.020] | 0.029*<br>[0.005, 0.055] | -0.001<br>[-0.005, 0.003] | Convergence Problems. |  | .355 | .048 |
| 2 | 1 | Intercept | 1.320*<br>[1.269, 1.371] | -0.029*<br>[-0.031, -0.027] | 0.135*<br>[0.121, 0.148] | NA | 13224.68 | .000 | .303 | .083 |
| 2 | 2 | Intercept | 1.321*<br>[1.269, 1.373] | -0.029*<br>[-0.032, -0.026] | 0.133*<br>[0.105, 0.160] | -0.001<br>[-0.004, 0.005] | 13226.65 | .000 | .303 | .083 |
| 2 | 3 | Intercept and DM offer | 1.314*<br>[1.266, 1.361] | -0.028*<br>[-0.031, -0.024] | 0.134*<br>[0.121, 0.147] | NA | 13180.36 | .000 | .304 | .078 |
| 2 | 4 | Intercept and DM offer | 1.314*<br>[1.266, 1.363] | -0.028*<br>[-0.032, -0.024] | 0.132*<br>[0.105, 0.160] | -0.001<br>[-0.004, 0.005] | 13182.34 | .004 | .304 | .078 |
| **2** | **5** | **Intercept, DM offer, and RC type** | **1.312\***<br>**[1.264, 1.359]** | **-0.028\***<br>**[-0.031, -0.024]** | **0.138\***<br>**[0.110, 0.165]** | **NA** | **13079.58** | **.999** | **.318** | **.079** |
| 2 | 6 | Intercept, DM offer, and RC type | 1.313*<br>[1.267, 1.359] | -0.028*<br>[-0.031, -0.025] | 0.135*<br>[0.099, 0.172] | -0.001<br>[-0.004, 0.005] | Convergence Problems. |  | .312 | .079 |

*Note.* AIC = Akaike Information Criterion. AICw = AIC weight. DM offer = Decision-maker offer. RC type = Receiver type (i.e., contextual-expectancy condition) dummy coded as 1 = Old Receiver (Context-expectant condition) and 0 = New Receiver (Context- not-expectant condition). Interaction = interaction effect between DM offer and RC type. Significant predictors that do not cross zero are marked with ‘\*’.

**Table S3.** Random effects structure, and confidence intervals thereof, for the best fitting model, across two studies.

| Experiment | Random Effect | Standard Deviation | Confidence Interval |  | Correlations |  |
| --- | --- | --- | --- | --- | --- | --- |
|  |  |  | 2.5% | 97.5% | Intercept | DM offer |
| 1 | Intercept | 0.221 | 0.182 | 0.27 |  |  |
| 1 | Decision-maker offer | 0.017 | 0.014 | 0.022 | -0.399 |  |
| 1 | Context-expectancy | 0.047 | 0.031 | 0.071 | -0.273 | 0.089 |
| 2 | Intercept | 0.172 | 0.14 | 0.211 |  |  |
| 2 | Decision-maker offer | 0.011 | 0.008 | 0.015 | 0.046 |  |
| 2 | Context-expectancy | 0.093 | 0.073 | 0.119 | -0.099 | 0.096 |

*Note.* DM offer = Decision-maker offer.

**Table S4.**
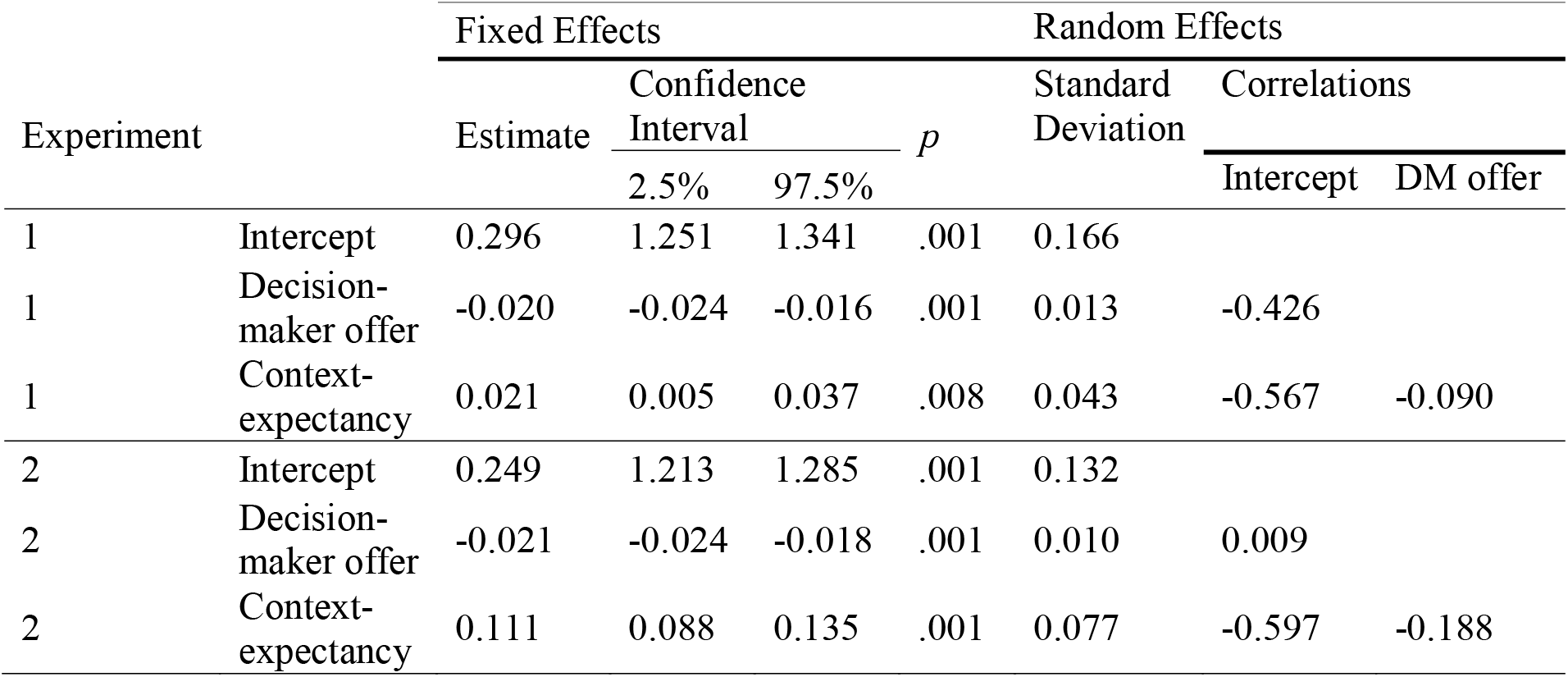
Fixed effects estimates and random effects structure for an alternative version of the winning model based on gaussian distribution with a log link function.

**Table S5.** Fixed effects estimates and random effects structure for an alternative version of the winning model based on gaussian distribution with an identity link function.

| Experiment |  | Fixed Effects |  |  |  | Random Effects |  |  |
| --- | --- | --- | --- | --- | --- | --- | --- | --- |
|  |  | Estimate | Confidence Interval |  | *p* | Standard Deviation | Correlations |  |
|  |  |  | 2.5% | 97.5% |  |  | Intercept | DM offer |
| 1 | Intercept | 1.362 | 1.302 | 1.421 | .001 | 0.219 |  |  |
| 1 | Decision-maker offer | -0.025 | -0.03 | -0.02 | .001 | 0.017 | -0.574 |  |
| 1 | Context-expectancy | 0.024 | 0.005 | 0.044 | .016 | 0.054 | -0.520 | -0.004 |
| 2 | Intercept | 1.301 | 1.255 | 1.347 | .001 | 0.169 |  |  |
| 2 | Decision-maker offer | -0.026 | -0.029 | -0.022 | .001 | 0.011 | -0.249 |  |
| 2 | Context-expectancy | 0.138 | 0.109 | 0.166 | .001 | 0.095 | -0.395 | -0.107 |

**Figure S1.**
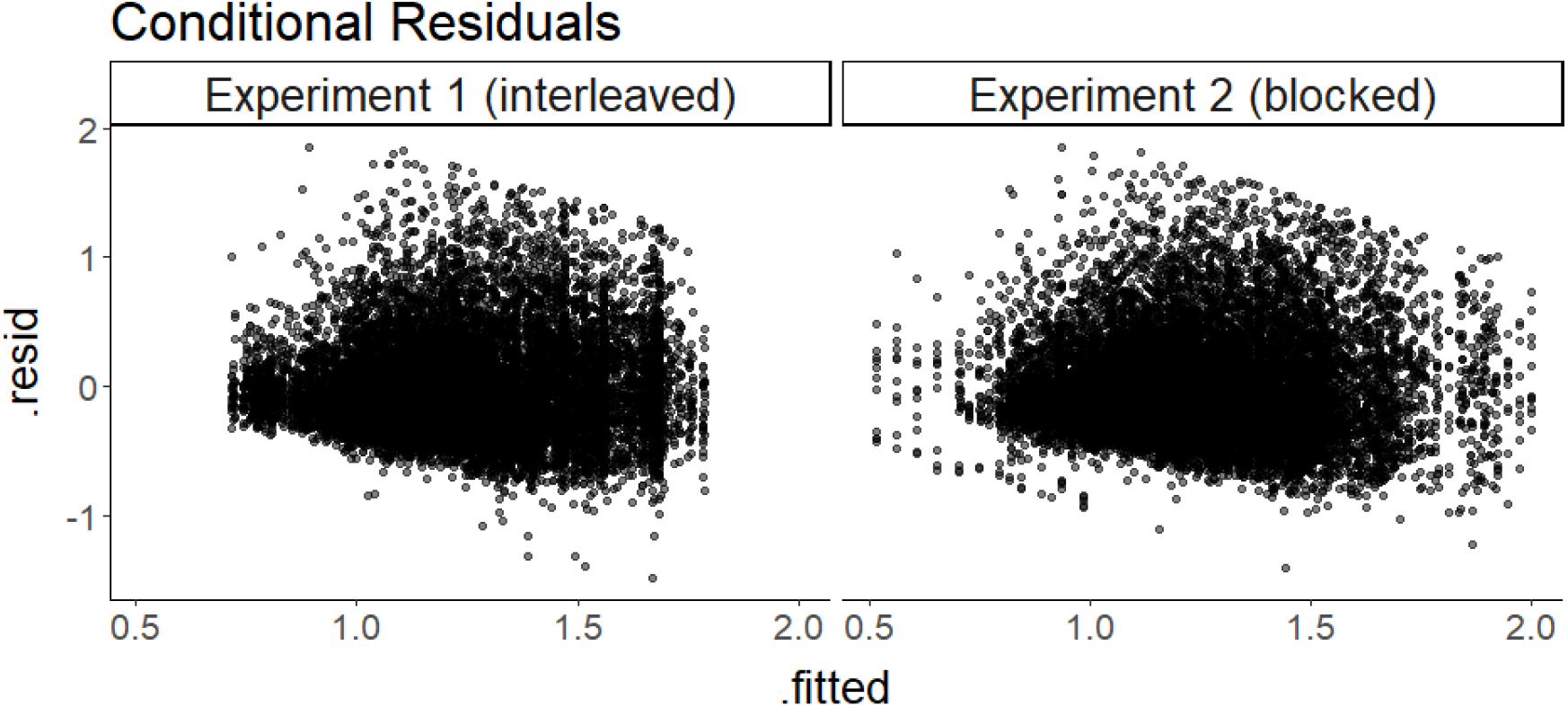
Diagnostic plot of model raw conditional residuals (y axis) by model predicted value (x axis). Note that increasing residual raw variance (heteroscedasticity) is expected for gamma models as the variance increases with the mean of the distribution.

**Figure S2.**
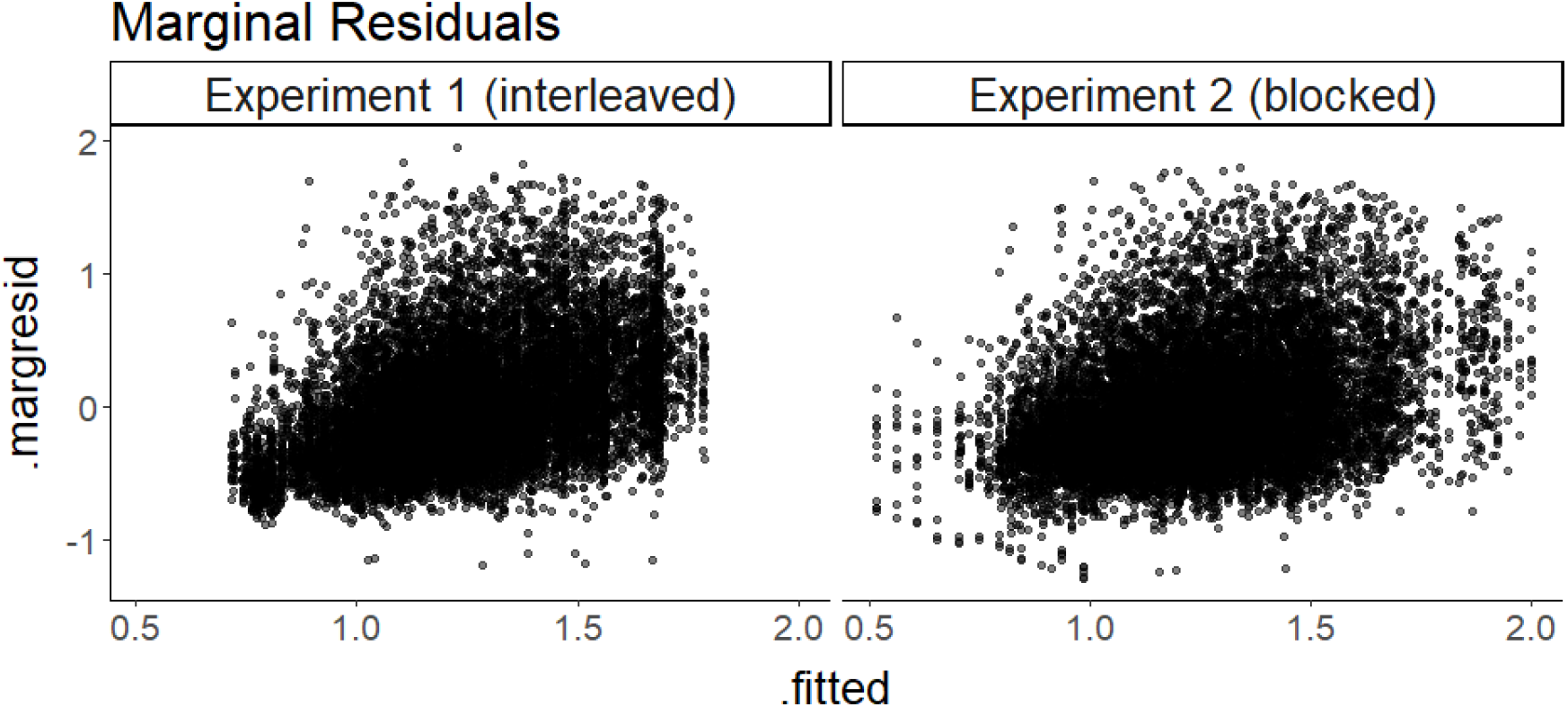
Diagnostic plot of model raw marginal residuals (y axis) by model predicted value (x axis). Note that increasing residual raw variance (heteroscedasticity) is expected for gamma models as the variance increases with the mean of the distribution.

**Figure S3.**
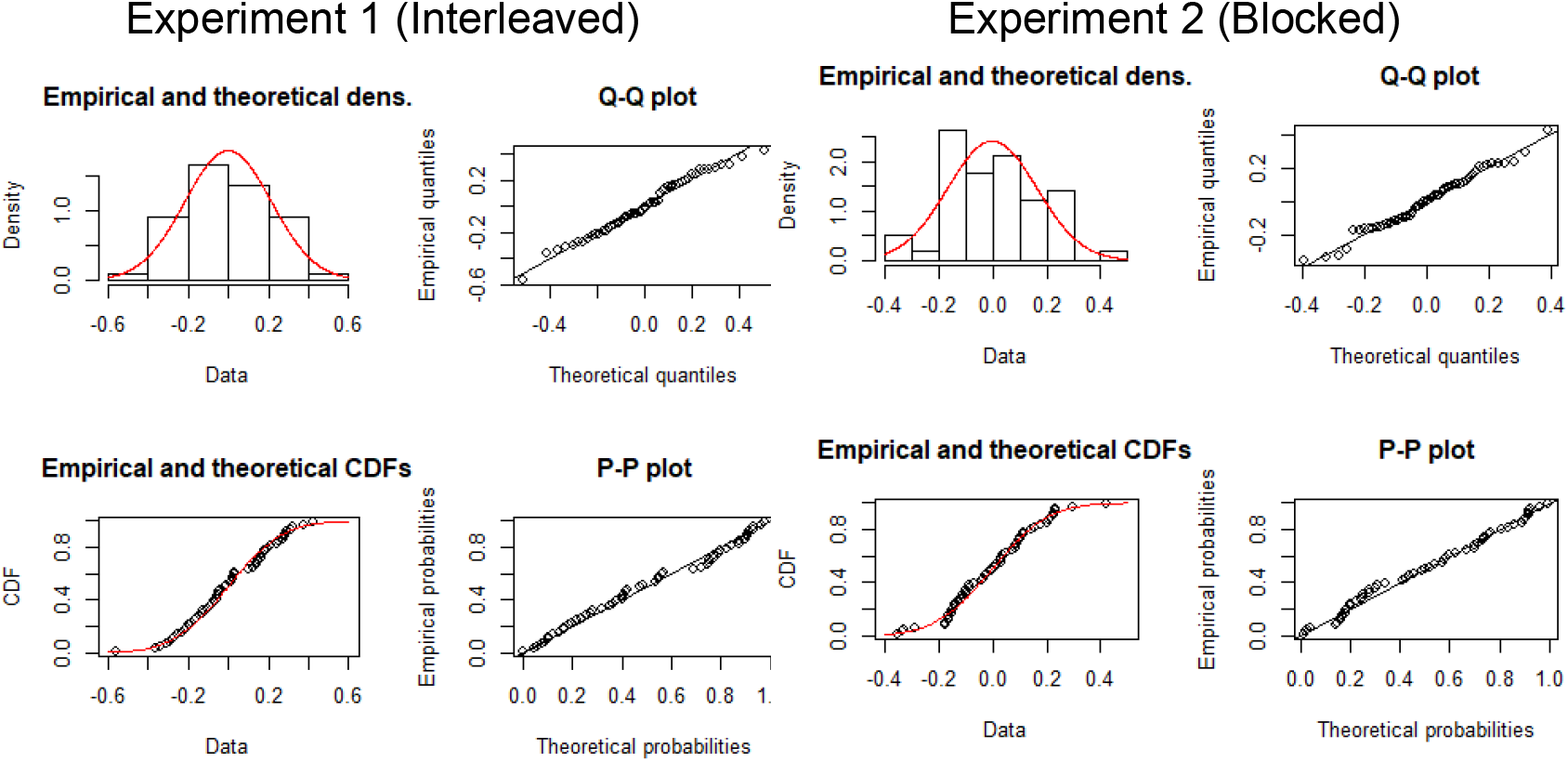
Diagnostic plots comparing the intercept random effect of the best-fitting RT model to a normal distribution, showing that the model assumptions are upheld.

**Figure S4.**
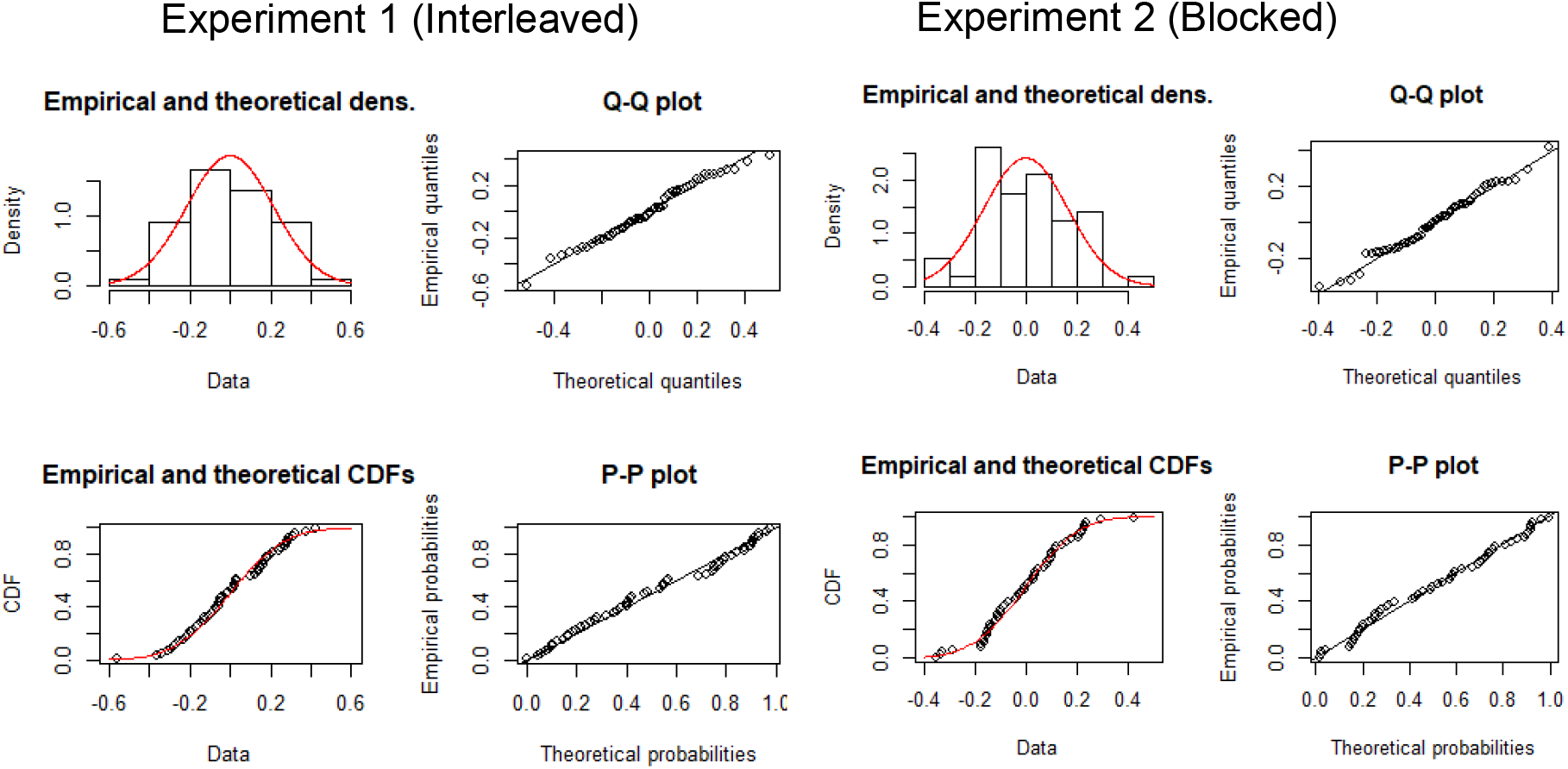
Diagnostic plots comparing the Decision-maker offer random effect of the best-fitting RT model to a normal distribution, showing that the model assumptions are upheld.

### Model Simulations

Figures S5–7 show how data simulated from our best-fitting model structure compare to the observed data at various levels of aggregation – experiment, context-expectancy, condition, and offer magnitude condition.

**Figure S5.**
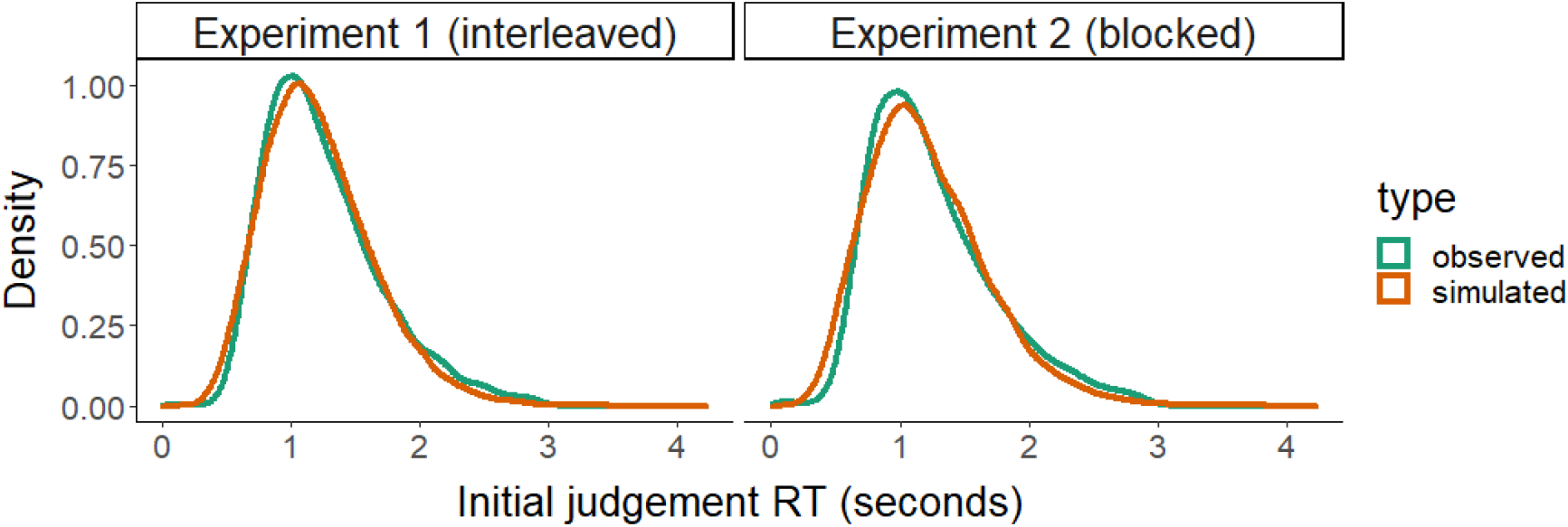
Comparison of observed RT data with data simulated from model structure at aggregate level.

**Figure S6.**
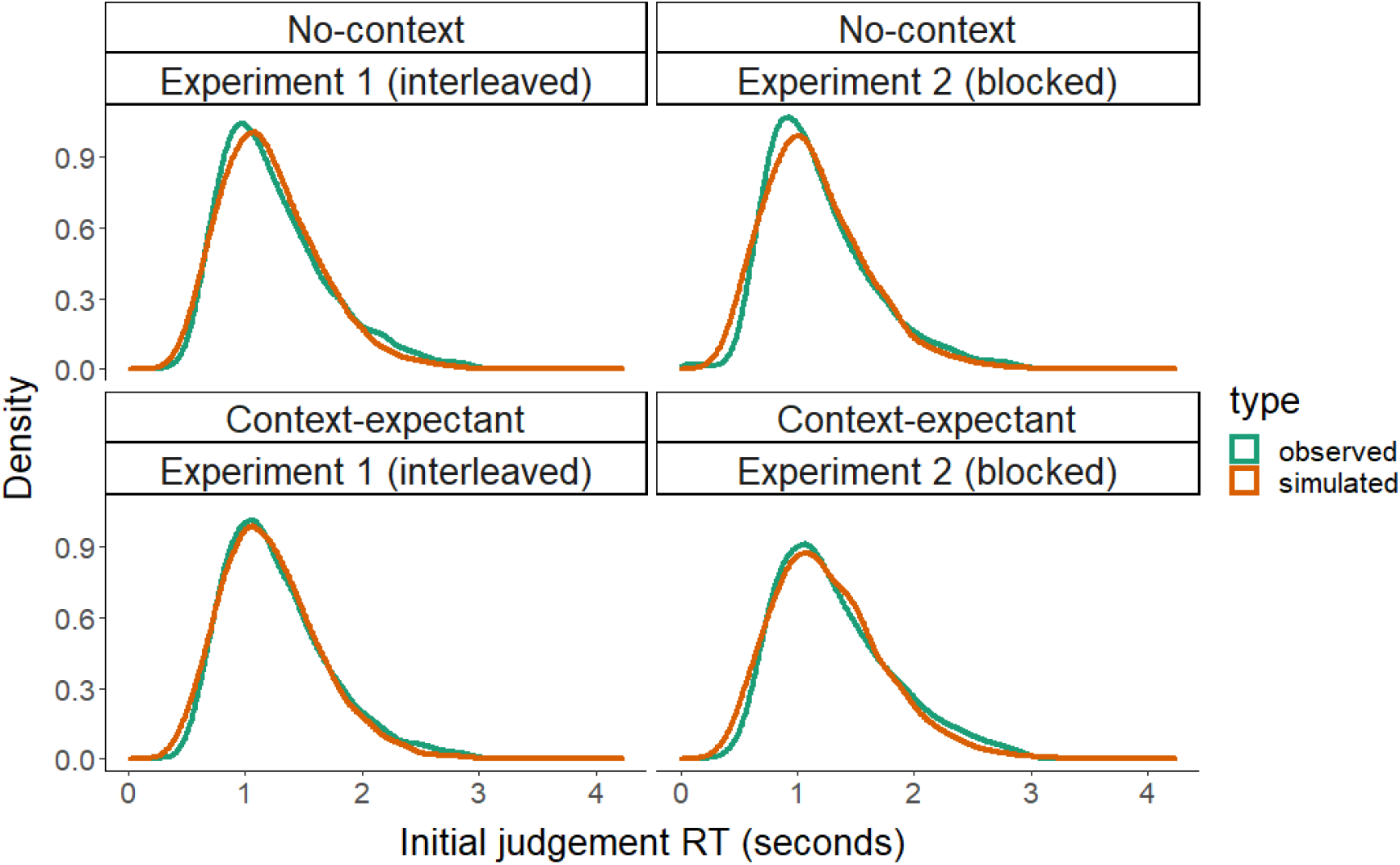
Comparison of observed RT data with data simulated from model structure, faceted by contextual-information condition.

**Figure S7.**
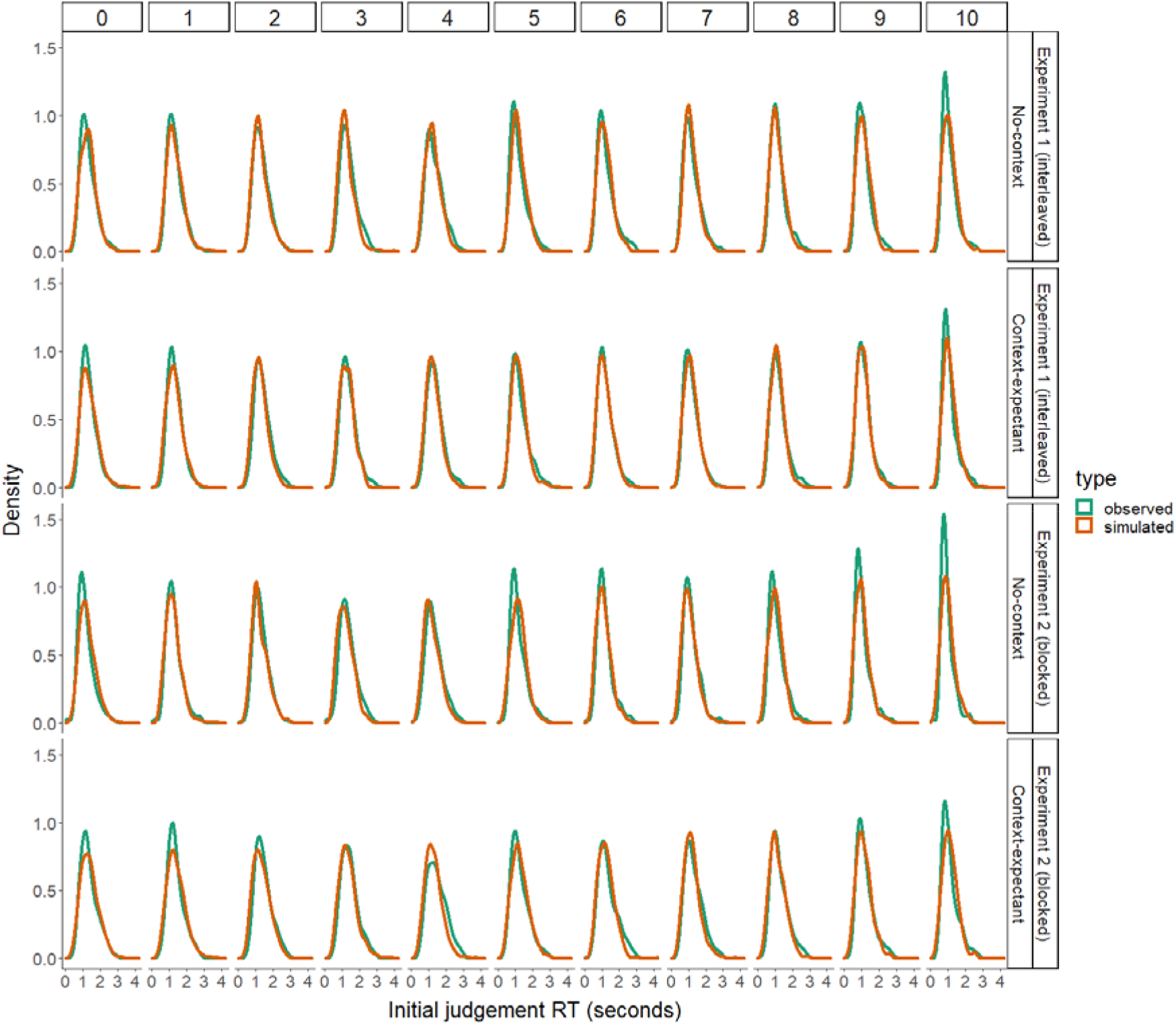
Comparison of observed RT data with data simulated from model structure, faceted by contextual-information condition and Decision-maker offer.

## Supplement 2: Diffusion Decision Model

### Model Simulations

Figures S8–10 show how data simulated from the two version of the fitted Diffusion Decision Models (DDM) using the Posterior Probability Check procedure compare to empirical data.

**Figure S8.**
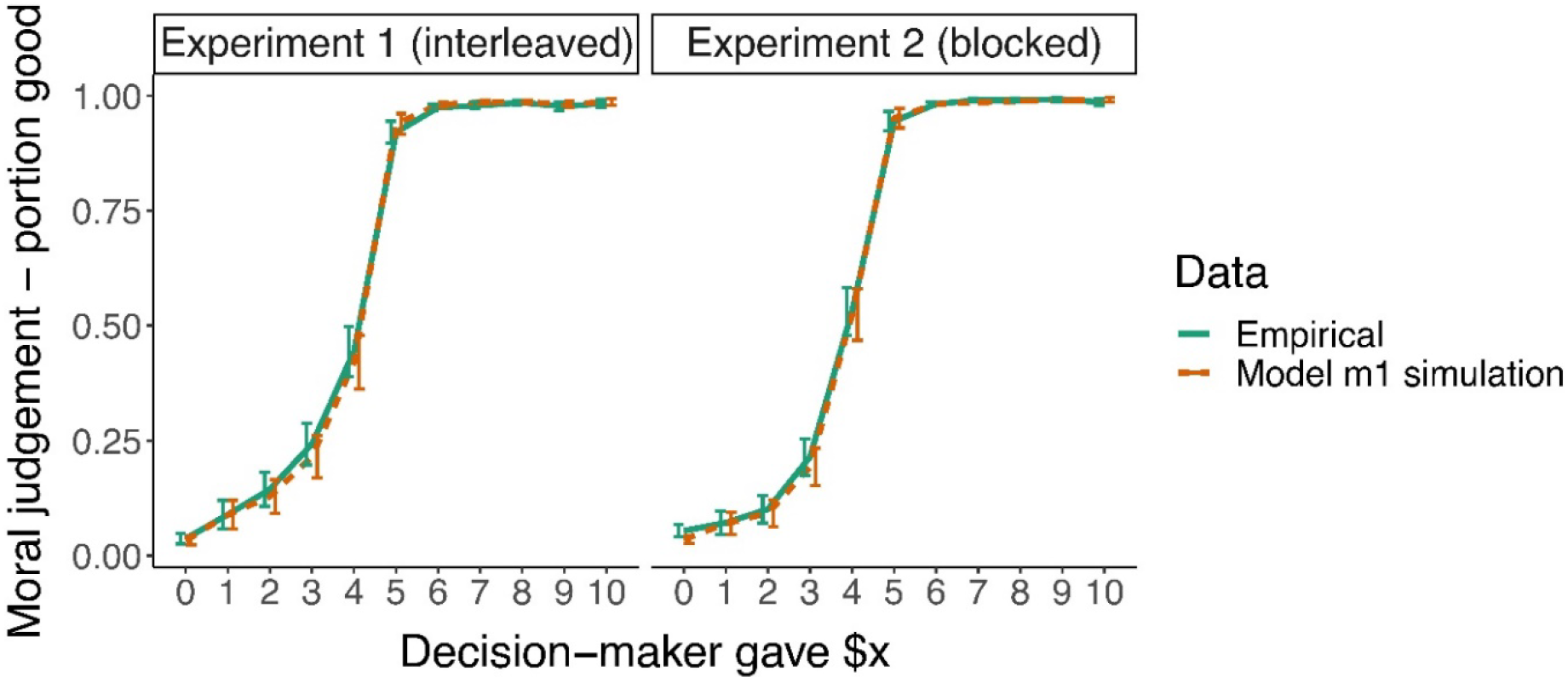
Comparison of empirical judgement data with data simulated by the PPC procedure from the hypothesised Diffusion Decision Model. Error bars depict the SEM.

**Figure S9.**
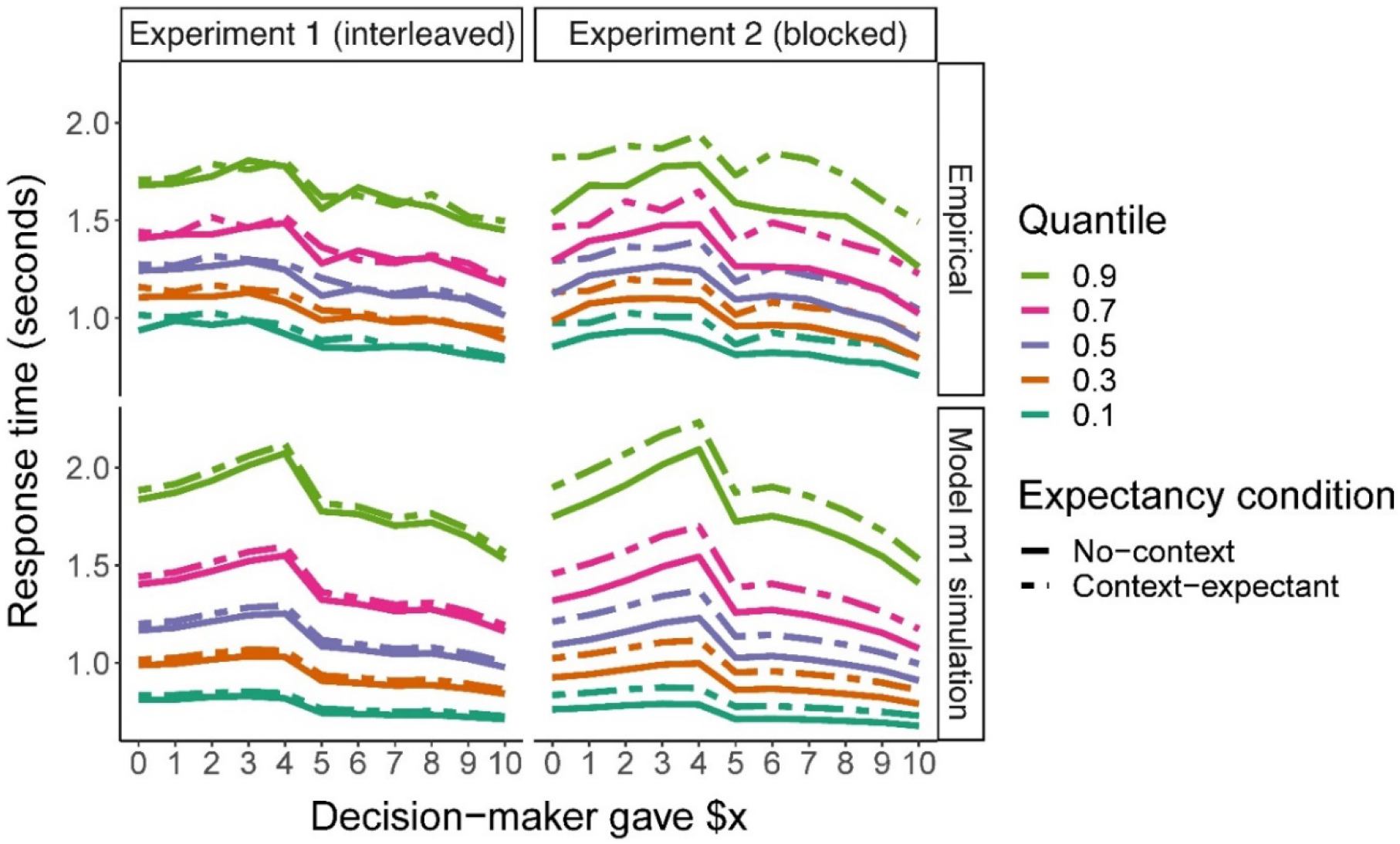
Comparison of empirical RT quantiles with data simulated by the PPC procedure from the hypothesised Diffusion Decision Model.

**Figure S10.**
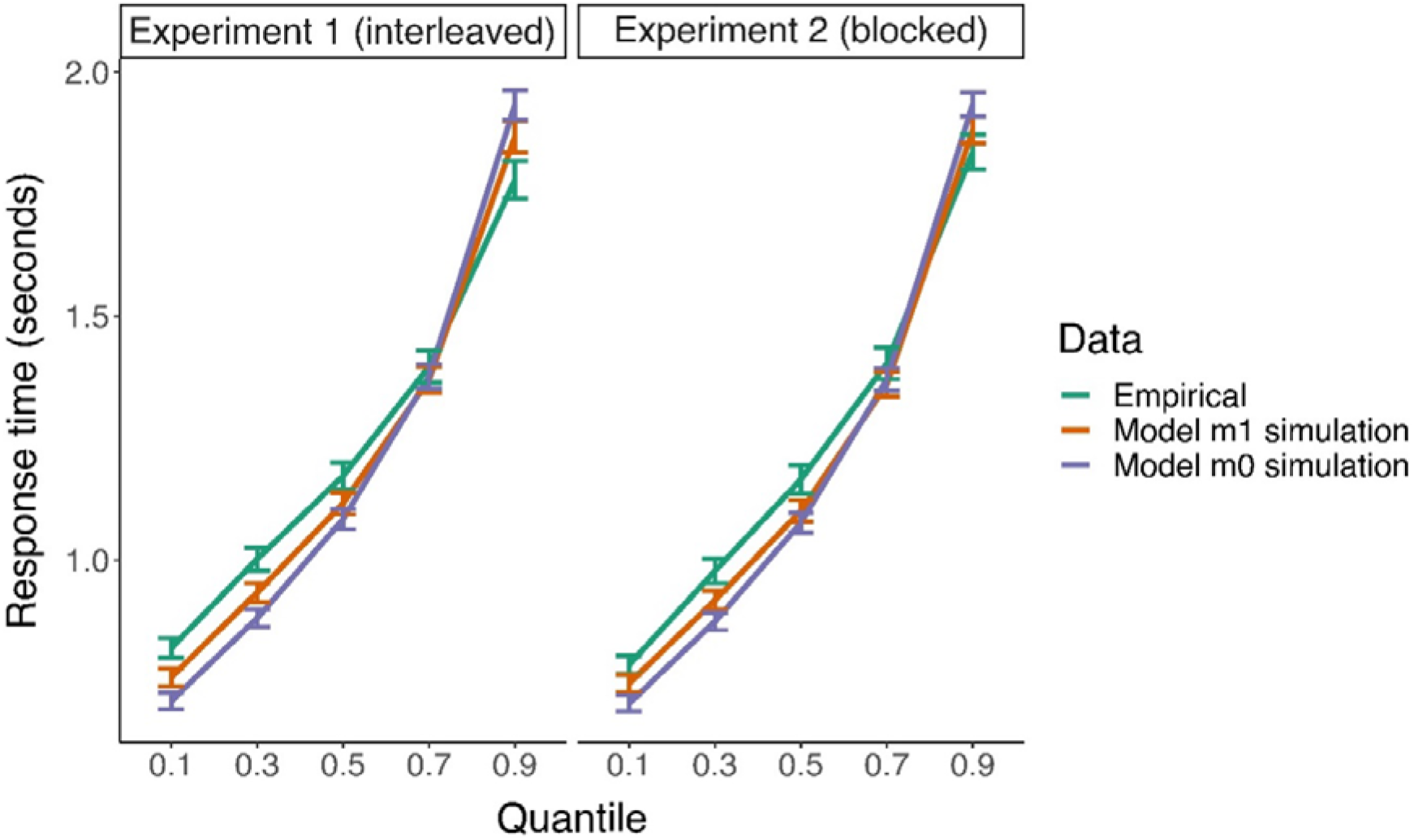
Comparison of the observed RT quantiles and RT quantiles from a dataset simulated by the PPC procedure from two fitted Diffusion Decision Models. Error bars depict the SEM. The hypothesised m1 model approximates the quantile structure of RT better than the null m0 model.

## References

1. Turiel, E. The Culture of Morality. (Cambridge University Press, 2001). doi:10.1017/CBO9780511613500

2. Miron, A. M., Warner, R. H. & Branscombe, N. R. Accounting for group differences in appraisals of social inequality: Differential injustice standards. Br. J. Soc. Psychol. 50, 342–353 (2011).

3. Sawaoka, T., Newheiser, A.-K. & Dovidio, J. F. Group-based biases in moral judgment: The role of shifting moral standards. Soc. Cogn. 32, 360–380 (2014).

4. Olson, J. G., McFerran, B., Morales, A. C. & Dahl, D. W. Wealth and welfare: Divergent moral reactions to ethical consumer choices. J. Consum. Res. 42, 879–896 (2016).

5. Haidt, J. & Baron, J. Social roles and the moral judgement of acts and omissions. Eur. J. Soc. Psychol. 26, 201–218 (1996).

6. Simpson, A., Laham, S. M. & Fiske, A. P. Wrongness in different relationships[: Relational context effects on moral judgment. J. Soc. Psychol. 156, 594–609 (2016).

7. Feather, N. T. & Deverson, N. H. Reactions to a motor-vehicle accident in relation to mitigating circumstances and the gender and moral worth of the driver. J. Appl. Soc. Psychol. 30, 77–95 (2000).

8. Feather, N. T. Judgments of deservingness: Studies in the psychology of justice and achievement. Personal. Soc. Psychol. Rev. 3, 86–107 (1999).

9. Andrejević, M., Feuerriegel, D., Turner, W., Laham, S. & Bode, S. Moral judgements of fairness-related actions are flexibly updated to account for contextual information. Sci. Rep. 10, 17828 (2020).

10. Shapiro, E. G. Effect of expectations of future interaction on reward allocations in dyads: Equity or equality. J. Pers. Soc. Psychol. 31, 873–880 (1975).

11. Deutsch, M. Equity, equality, and need: What determines which value will be used as the basis of distributive justice? J. Soc. Issues 31, 137–149 (1975).

12. Hysom, S. J. & Fişek, M. H. Situational determinants of reward allocation: The equity-equality equilibrium model. Soc. Sci. Res. 40, 1263–1285 (2011).

13. Diekmann, A. The power of reciprocity: Fairness, reciprocity, and stakes in variants of the dictator game. J. Conflict Resolut. 48, 487–505 (2004).

14. Fehr, E. & Fischbacher, U. Third-party punishment and social norms. Evol. Hum. Behav. 25, 63–87 (2004).

15. Nowak, M. A. & Sigmund, K. Evolution of indirect reciprocity by image scoring. Nature 393, 573–577 (1998).

16. Meristo, M. & Surian, L. Do infants detect indirect reciprocity? Cognition 129, 102–113 (2013).

17. Adams, J. S. Inequity in social exchange. in Advances in Experimental Social Psychology (ed. Berkowitz, L.) 2, 267–299 (Academic Press, 1965).

18. Messick, D. M. & Schell, T. Evidence for an equality heuristic in social decision making. Acta Psychol. (Amst). 80, 311–323 (1992).

19. Fehr, E. & Schmidt, K. M. A theory of fairness, competition, and cooperation. Q. J. Econ. 114, 817–868 (1999).

20. Forstmann, B. U. et al. Cortico-striatal connections predict control over speed and accuracy in perceptual decision making. Proc. Natl. Acad. Sci. U. S. A. 107, 15916–15920 (2010).

21. Voss, A., Rothermund, K. & Voss, J. Interpreting the parameters of the diffusion model: An empirical validation. Mem. Cogn. 32, 1206–1220 (2004).

22. Green, N., Biele, G. P. & Heekeren, H. R. Changes in neural connectivity underlie decision threshold modulation for reward maximization. J. Neurosci. 32, 14942–14950 (2012).

23. Starns, J. J. & Ratcliff, R. Age-related differences in diffusion model boundary optimality with both trial-limited and time-limited tasks. Psychon. Bull. Rev. 19, 139–145 (2012).

24. Dunovan, K. & Verstynen, T. Errors in action timing and inhibition facilitate learning by tuning distinct mechanisms in the underlying decision process. J. Neurosci. 39, 2251–2264 (2019).

25. Bond, K., Dunovan, K. & Verstynen, T. Value-conflict and volatility influence distinct decision-making processes. in (2019). doi:10.32470/ccn.2018.1068-0

26. Summerfield, C. & Koechlin, E. Economic value biases uncertain perceptual choices in the parietal and prefrontal cortices. Front. Hum. Neurosci. 4, 1–12 (2010).

27. Monroe, A. E. & Malle, B. F. People systematically update moral judgments of blame. J. Pers. Soc. Psychol. 116, 215–236 (2019).

28. Siegel, J. Z., Mathys, C., Rutledge, R. B. & Crockett, M. J. Beliefs about bad people are volatile. Nat. Hum. Behav. 2, 750–756 (2018).

29. Abele, A. Thinking about thinking: Causal, evaluative and finalistic cognitions about social situations. Eur. J. Soc. Psychol. 15, 315–332 (1985).

30. Fiske, S. T. Attention and weight in person perception: The impact of negative and extreme behavior. J. Pers. Soc. Psychol. 38, 889–906 (1980).

31. Pratto, F. & John, O. P. Automatic vigilance[: The attention-grabbing power of approach- and avoidance-related social information. J. Personal. Soc. Psychol. 61, 380–391 (1991).

32. Baumeister, R. F., Bratslavsky, E., Finkenauer, C. & Vohs, K. D. Bad is stronger than good. Rev. Gen. Psychol. 5, 323–370 (2001).

33. Ratcliff, R. A theory of memory retrieval. Psychol. Rev. 85, 59–108 (1978).

34. Smith, P. L. & Ratcliff, R. Psychology and neurobiology of simple decisions. Trends in Neuroscience 27, 161–168 (2004).

35. Ratcliff, R., Smith, P. L., Brown, S. D. & McKoon, G. Diffusion Decision Model: Current issues and history. Trends Cogn. Sci. 20, 260–281 (2016).

36. Ratcliff, R., Gomez, P. & McKoon, G. A diffusion model account of the lexical decision task. Psychol. Rev. 111, 159–182 (2004).

37. Gomez, P., Ratcliff, R. & Perea, M. A model of the go/no-go task. J. Exp. Psychol. Gen. 136, 389–413 (2007).

38. Kiani, R. & Shadlen, M. N. Representation of confidence associated with a decision by neurons in the parietal cortex. Science 324, 759–764 (2009).

39. Ratcliff, R., Thapar, A. & McKoon, G. A diffusion model analysis of the effects of aging on recognition memory. J. Mem. Lang. 50, 408–424 (2004).

40. Cook, E. P. & Maunsell, J. H. R. Dynamics of neuronal responses in macaque MT and VIP during motion detection. Nat. Neurosci. 5, 985–994 (2002).

41. Gold, J. I. & Shadlen, M. N. The neural basis of decision making. Annual Review of Neuroscience 30, 535–574 (2007).

42. Milosavljevic, M., Malmaud, J., Huth, A., Koch, C. & Rangel, A. The Drift Diffusion Model can account for the accuracy and reaction time of value-based choices under high and low time pressure. Judgm. Decis. Mak. 5, 437–449 (2010).

43. Gallotti, R. & Grujić, J. A quantitative description of the transition between intuitive altruism and rational deliberation in iterated Prisoner’s Dilemma experiments. Sci. Rep. 9, 1–11 (2019).

44. Hutcherson, C. A., Bushong, B. & Rangel, A. A neurocomputational model of altruistic choice and its implications. Neuron 87, 451–463 (2015).

45. Son, J. Y., Bhandari, A. & Feldman Hall, O. Crowdsourcing punishment: Individuals reference group preferences to inform their own punitive decisions. Sci. Rep. 9, 1–15 (2019).

46. Pärnamets, P. et al. Biasing moral decisions by exploiting the dynamics of eye gaze. Proc. Natl. Acad. Sci. U. S. A. 112, 4170–4175 (2015).

47. Plassmann, H., O’Doherty, J. & Rangel, A. Orbitofrontal cortex encodes willingness to pay in everyday economic transactions. J. Neurosci. 27, 9984–9988 (2007).

48. Basten, U., Biele, G., Heekeren, H. R. & Fiebach, C. J. How the brain integrates costs and benefits during decision making. Proc. Natl. Acad. Sci. 107, 21767–21772 (2010).

49. Andrejević, M. et al. How do basic personality traits map onto moral judgements of fairness-related actions? PsyArXiv (2020). Preprint at https://doi.org/10.31234/osf.io/e3uxb

50. Spiegelhalter, D. J., Best, N. G., Carlin, B. P. & Van Der Linde, A. Bayesian measures of model complexity and fit. J. R. Stat. Soc. Ser. B Stat. Methodol. 64, 583–616 (2002).

51. Pärnamets, P., Richardson, D. & Balkenius, C. Modelling moral choice as a diffusion process dependent on visual fixations. Proc. Annu. Meet. Cogn. Sci. Soc. (2014).

52. Pärnamets, P. et al. Changing minds by tracking eyes: Dynamical systems, gaze and moral decisions. Proc. Annu. Meet. Cogn. Sci. Soc. 1115–1120 (2013).

53. Ditterich, J. Stochastic models of decisions about motion direction: Behavior and physiology. Neural Networks 19, 981–1012 (2006).

54. Churchland, A. K., Kiani, R. & Shadlen, M. N. Decision-making with multiple alternatives. Nat. Neurosci. 11, 693–702 (2008).

55. Ratcliff, R. & Rouder, J. N. Modeling response times for two-choice decisions. Psychological Science 9, (1998).

56. Ratcliff, R. Parameter variability and distributional assumptions in the diffusion model. Psychol. Rev. 120, 281–292 (2013).

57. Kleiner, M. et al. What’s new in Psychtoolbox-3? Perception (2007). doi:10.1068/v070821

58. Güth, W., Schmittberger, R. & Schwarze, B. An experimental analysis of ultimatum bargaining. J. Econ. Behav. Organ. 3, 367–388 (1982).

59. Lee, K. & Ashton, M. C. Psychometric properties of the HEXACO-100. Assessment 25, 543–556 (2018).

60. Graham, J., Haidt, J. & Nosek, B. A. Liberals and conservatives rely on different sets of moral foundations. J. Pers. Soc. Psychol. 96, 1029–1046 (2009).

61. Ho, A. K. et al. The nature of social dominance orientation: Theorizing and measuring preferences for intergroup inequality using the new SDO□ scale. J. Pers. Soc. Psychol. 109, 1003–1028 (2015).

62. Piazza, J. & Sousa, P. Religiosity, political orientation, and consequentialist moral thinking. Soc. Psychol. Personal. Sci. 5, 334–342 (2013).

63. Lo, S. & Andrews, S. To transform or not to transform: using generalized linear mixed models to analyse reaction time data. Front. Psychol. 6, 1–16 (2015).

64. De Boeck, P. & Jeon, M. An overview of models for response times and processes in cognitive tests. Front. Psychol. 10, (2019).

65. Maris, E. Additive and multiplicative models for gamma distributed random variables, and their application as psychometric models for response times. Psychometrika 58, 445–469 (1993).

66. Palmer, E. M., Horowitz, T. S., Torralba, A. & Wolfe, J. M. What are the shapes of response time distributions in visual search? J. Exp. Psychol. Hum. Percept. Perform. 37, 58–71 (2011).

67. Zandt, T. Van. Analysis of response time distributions. Stevens’ Handbook of Experimental Psychology (2002). doi:doi:10.1002/0471214426.pas0412

68. Brooks, M. E. et al. glmmTMB balances speed and flexibility among packages for zero-inflated generalized linear mixed modeling. R J. 9, 378–400 (2017).

69. Akaike, H. A new look at the statistical model identification. IEEE Trans. Automat. Contr. 19, 716–723 (1974).

70. Schwarz, G. Estimating the dimension of a model. Ann. Stat. 6, 461–464 (1978).

71. Vrieze, S. I. Model selection and psychological theory: A discussion of the differences between the Akaike information criterion (AIC) and the Bayesian information criterion (BIC). Psychol. Methods 17, 228–243 (2012).

72. Wagenmakers, E. J. & Farrell, S. AIC model selection using Akaike weights. Psychon. Bull. Rev. 11, 192–196 (2004).

73. Bolker, B. M. et al. Generalized linear mixed models: a practical guide for ecology and evolution. Trends Ecol. Evol. 24, 127–135 (2009).

74. Meade, B., Lafayette, L., Sauter, G. & Tosello, D. Spartan HPC-Cloud hybrid: Delivering performance and flexibility. (2017). doi:10.4225/49/58ead90dceaaa

75. Wiecki, T. V., Sofer, I. & Frank, M. J. HDDM: Hierarchical bayesian estimation of the drift-diffusion model in Python. Front. Neuroinform. 7, 1–10 (2013).

76. Ratcliff, R. Estimating parameters of the diffusion model: Approaches to dealing with contaminant reaction times and parameter variability. 9, 438–481 (2002).

